# Eco-evolutionary dynamics of massive, parallel bacteriophage outbreaks in compost communities

**DOI:** 10.1101/2023.07.31.550844

**Authors:** Jeroen Meijer, Petros Skiadas, Paul B. Rainey, Paulien Hogeweg, Bas E. Dutilh

## Abstract

Bacteriophages are important drivers of microbial ecosystems, yet their dynamics in complex natural communities remain poorly understood compared to simplified laboratory systems. To address this gap, we analyzed viral dynamics across 20 compost-derived microbial communities propagated for one year in mesocosms. Communities clustered into two distinct types, each dominated by different cellulose-degraders and comprising hundreds of genera. In one type, we observed massive, parallel outbreaks of Theomophage, a previously undescribed bacteriophage, reaching 74% of metagenomic sequencing reads, the largest bacteriophage outbreak documented to date. Theomophage populations in isolated communities were composed of a single genotype that showed striking evolutionary stability throughout the experiment. In contrast, following experimental viral migration between mesocosms, all Theomophage populations showed rapid evolution via recombination between preexisting genotypes, replacement of ancestral lineages, and, upon successful migration to mesocosms of the alternate community type where Theomophage was initially absent, rapid acquisition of novel mutations that swept local populations and then spread to other mesocosms. Our study reveals the spatial and temporal scales at which bacteriophage microdiversity evolves in complex communities. It further shows that mixing of viral communities — likely common in natural systems — can rapidly accelerate bacteriophage evolution.

## Introduction

Viruses that infect bacteria (bacteriophages or phages) are ubiquitous and ecologically influential components of all microbial ecosystems, from oceans and soils to host-associated microbiomes. They regulate bacterial populations, drive nutrient cycling, and shape bacterial evolution by facilitating horizontal gene transfer^1^. Phages may also shape community structure and function, though evidence is mixed^2^. These ecological and evolutionary effects arise from highly specific interactions between phages and their bacterial hosts, which are typically shaped by antagonistic coevolution^3–6^. Identifying the biotic and abiotic factors that govern phage eco-evolutionary dynamics is essential for predicting microbial community diversity, stability, and function, and for guiding applied interventions such as bacteriophage therapy^7^.

Current knowledge of phage ecology and evolution is largely shaped by two contrasting study designs: simplified *in vitro* experimental systems in nutrient rich media, typically consisting of one or a few phage-host pairs, and observational surveys of complex natural ecosystems using metagenomics or large-scale isolation and phenotyping^8–10^. Experiments with single phage-host pairs show coevolution proceeding via series of selective sweeps, in which bacterial receptor modification or loss confers resistance to infection, countered by compensatory changes in phage tail proteins^4,5,11^. In contrast, studies of natural ecosystems showed that bacterial resistance is shaped more by variation in defense systems than by receptor evolution^3,12,13^, and often reveal patterns of microdiversity suggesting long-term coexistence of resistant and susceptible hosts^14,15^. These differences indicate that selective pressures in simplified laboratory conditions differ fundamentally from those in complex natural environments. More broadly, our ability to extrapolate from laboratory models to predict viral dynamics in complex environments remains limited^9,10,16^, prompting calls for experimental models that incorporate greater ecological complexity while retaining experimental tractability^8–10,17^.

Several features of natural ecosystems likely contribute to these differences^9,10,17^. Natural communities harbor extensive macro-(taxonomic) and micro-(strain-level) diversity, in contrast to the isogenic populations typical of lab experiments. In diverse communities, genetic novelty arises not only through mutation, but also recombination and horizontal gene transfer (HGT). Increasing macrodiversity beyond a single phage-host pair can change both ecological and evolutionary trajectories (recently reviewed in ^9,17^). Natural ecosystems are also spatially heterogeneous, comprising fragmented patches (e.g. skin pores, gut crypts, marine snow, soil particles) connected by restricted dispersal, which produces local populations linked intermittently by migration^18^. Such metapopulation dynamics are inherently different from those in homogeneous laboratory systems^19–21^.

Microbial communities derived from natural ecosystems and propagated under laboratory conditions provide a promising middle ground between simplified lab systems and the full complexity of natural environments^22–27^. When grown on complex carbohydrates they maintain high taxonomic and functional diversity yet are amenable to longitudinal sampling and controlled perturbation. These systems have been used to investigate bacterial community assembly^22–26^, function^24^, and strain-level interactions^27^, but applications to viruses remain limited^28,29^. Here, using compost-derived microbial communities grown on paper, we investigate how standing microdiversity, bacteriophage migration, and community context shape bacteriophage eco-evolutionary dynamics. By integrating genotype-resolved viromics with a tractable yet complex system, we show that bacteriophage evolution can range from stasis to rapid change, depending on opportunities for migration.

## Results

### Compost communities cluster in two distinct and stable community types

To track the eco-evolutionary dynamics of bacteriophages in a complex system at high temporal resolution, we analyzed 184 shotgun metagenomes from a long-term mesocosm experiment with compost communities^30^. For clarity, we summarize the experimental setup. Ten independent founding communities were established by sampling 1 g of garden compost from different locations on a compost heap. Each community was incubated in mesocosms containing 20 mL nitrogen-limited minimal M9 medium supplemented with cellulose (4 cm^2^ paper) as the sole carbon source (**Fig.1a**). Following two weeks of incubation, 1 mL slurry was transferred to 19 mL fresh medium and cellulose paper, then incubated for another two weeks. After this acclimatization period (week 0), two serial transfer regimes were created from each mesocosm leading to 2×10 paired communities. In the closed regime, every 2 weeks 1 mL of cellulosic slurry was transferred to 19 mL fresh medium with cellulose paper. In the open regime, transfers were conducted identically, except that at every transfer, 1 mL of 0.2 μm-filtered viral fractions (hereafter the Mobile Genetic Element, MGE cocktail) were collected, pooled, and redistributed among all 10 open mesocosms. This design allowed migration of bacteriophages and other mobile genetic elements between open mesocosms, repeatedly exposing them to new hosts and potentially affecting their eco-evolutionary dynamics. Shotgun sequencing was performed at weeks 0, 2, 4, 6, 8, 20, 32, 40, and 48 for both regimes, with additional sequencing of the MGE cocktail at weeks 2, 4, 6, and 8. This enabled us to track viral migration via the MGE cocktail, and compare cellular and viral dynamics in closed mesocosms with their paired open counterparts.

**Figure 1.**
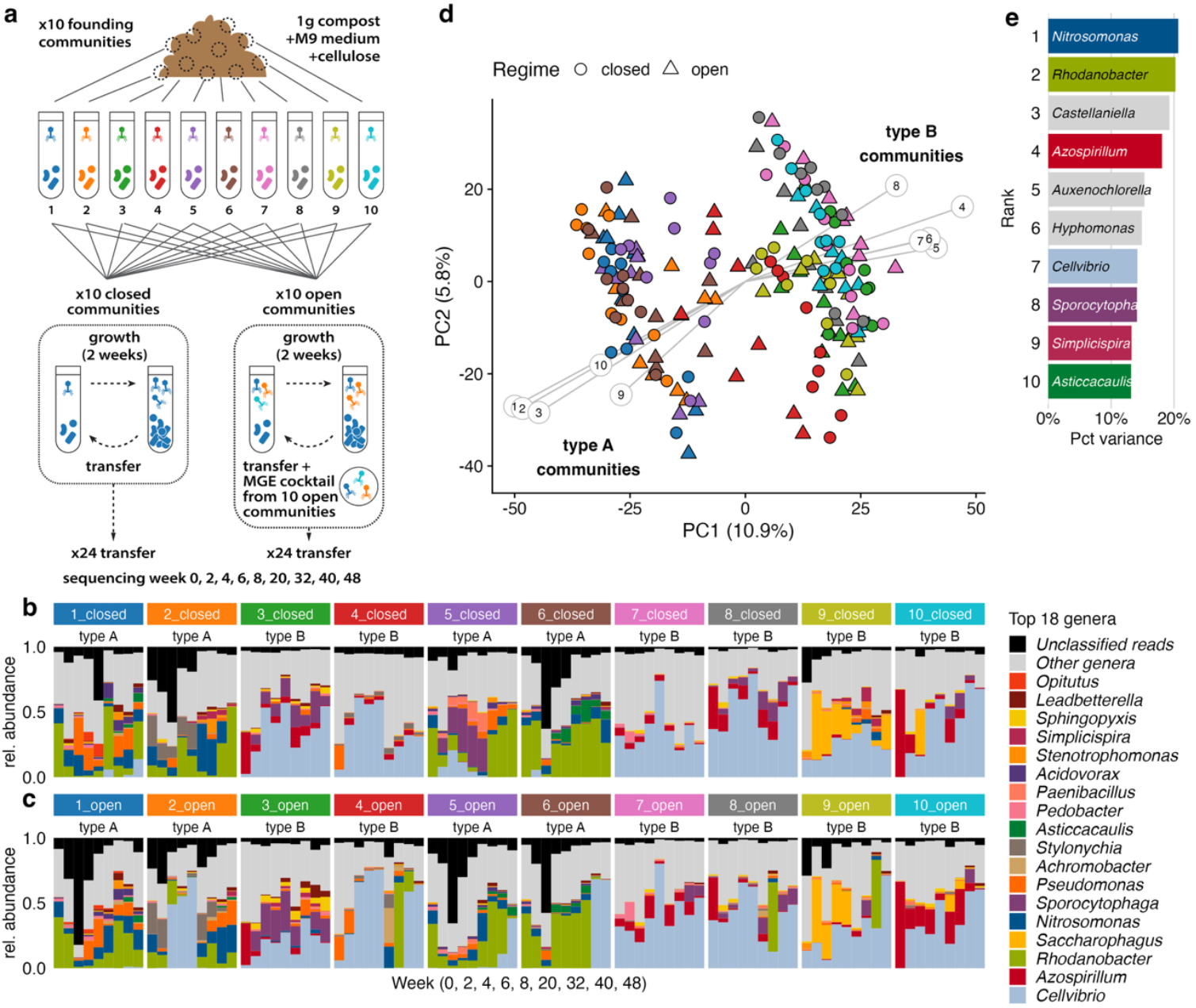
Compost communities cluster in two stable, distinct community types. **a**. Experimental design of Quistad et al^30^. Ten founding communities were established from garden compost and propagated on cellulose paper by serial transfer (2 weeks) for 48 weeks. In the closed regime communities were propagated in isolation. In the open regime, a mobile genetic element cocktail was added at every transfer, consisting of pooled, 0.2 μm-filtered material from all open communities (Methods). **b** Relative abundance (fraction of mapped metagenome reads) of the 18 most abundant genera in closed mesocosms (annotated with RAT). Unclassified reads (black) often represent viral reads (see **Fig.2**). **c** same as (b), for open mesocosms. **d** Principal component analysis (PCA) of centered log-ratio transformed genus-level relative abundances reveals two distinct community types along PC1: type A, rhodanobacter-dominated and type B, cellvibrio-dominated. Community type of mesocosms is stable over time and consistent across paired closed and open mesocosms, suggesting a strong founder effect. Grey lines indicate the top 10 genera contributing most to PC1 and PC2, numbered as in (e). For separate PCAs of closed and open regimes see Fig.S8. **e** Proportion of variance in PC1 and PC2 explained by the top 10 contributing genera. Numbers correspond to (d) and colors match (b) and (c).

We began by assembling the metagenomes and characterizing community composition, to provide context for viral eco-evolutionary dynamics. Taxonomic annotation of contigs and bins was performed using CAT^31^ and taxon relative abundances were calculated with RAT^32^ (**Fig.1b**, median 96.1% of sample reads mapped, **Fig.S1-2**; Methods). Consistent with previous studies^25,30^, this revealed diverse communities with mesocosms containing on average 150.3 ± 36.9 genera (mean ± standard deviation, mean 58.2% of sample reads annotated at genus rank, **Table S1; Fig.S1**). Principal component analysis (PCA) of genus-rank community composition revealed two distinct community types separated along the first component (**Fig.1bd**). Mesocosms closed_1, closed_2, closed_5, and closed_6 were dominated by genera including *Rhodanobacter* and *Nitrosomonas* (hereafter community type A), whereas the remaining 6 closed mesocosms were enriched in *Cellvibrio* and *Azospirillium* (community type B; **Fig.S4-5**). These community types were also apparent at higher taxonomic ranks and remained stable over 48 weeks, with dynamics primarily along the second PC (**Fig.S6-7**).

The two distinct community types were consistent across all paired closed and open mesocosms, suggesting a strong founder effect (**Fig.1cd**). This may reflect local sub-communities already present in different regions of the compost heap at the time of sampling, or alternatively, two distinct ecological attractors that emerged during the acclimatization period preceding the first sequenced time point. Stochastic diversity loss during acclimatization of natural communities to lab conditions can result in divergent communities and multistability^24–26^. Biopolymer degrading communities are structured by hierarchical cross-feeding and substrate preferences, with different degraders producing distinct metabolite profiles that influence downstream community assembly of specialized byproduct consumers^23^. We hypothesize that community type A and B are structured around different cellulose degraders, as *Cellvibrio* and *Rhodanobacter* are both associated with cellulose degradation and MAGs of these genera assembled from this dataset contain cellulases^33–35^. However, since cellulases are a highly heterogeneous enzyme group and their predicted genomic presence does not always indicate cellulolytic activity^36^, this remains to be experimentally validated.

### Massive, parallel outbreaks of a novel *Schitovirus* in type A communities

To investigate the viral dynamics in the compost mesocosms, we screened the assembled metagenomes using WhatThePhage^37^, a pipeline that integrates twelve virus identification tools (Methods). This analysis identified 140,378 predicted viral contigs, including 33 complete genomes, 131 high-quality (>90% complete), and 234 medium-quality genomes (50-90% complete, estimated with CheckV^38^). Among these, one 63,535 bp viral contig (estimated 100% complete) stood out due to its high abundance. Further analysis identified this contig as a representative of a new subfamily in the *Schitoviridae* bacteriophage family, which we named Theomophage after its place of origin, Place Théodore-Monod in Paris, France^30^ (see section “Theomophage represents a new *Schitoviridae* subfamily”). Between weeks 4-8 of the experiment, massive outbreaks of Theomophage occurred in all closed and open type-A mesocosms, with no outbreak detected in any type-B mesocosm (**Fig.2a**). These outbreaks reached abundances of up to 74.3% of total community reads (**Fig.2a**). Notably, outbreaks in the open mesocosms were often larger or occurred 2-6 weeks earlier than in their paired closed mesocosm (**Fig.2**), suggesting that Theomophage migration between open mesocosms accelerated and intensified outbreaks. Note that unclassified reads, including Theomophage reads, were not included in the PCA that delineated type-A and type-B communities, so Theomophage outbreaks did not drive the separation between community types.

**Figure 2.**
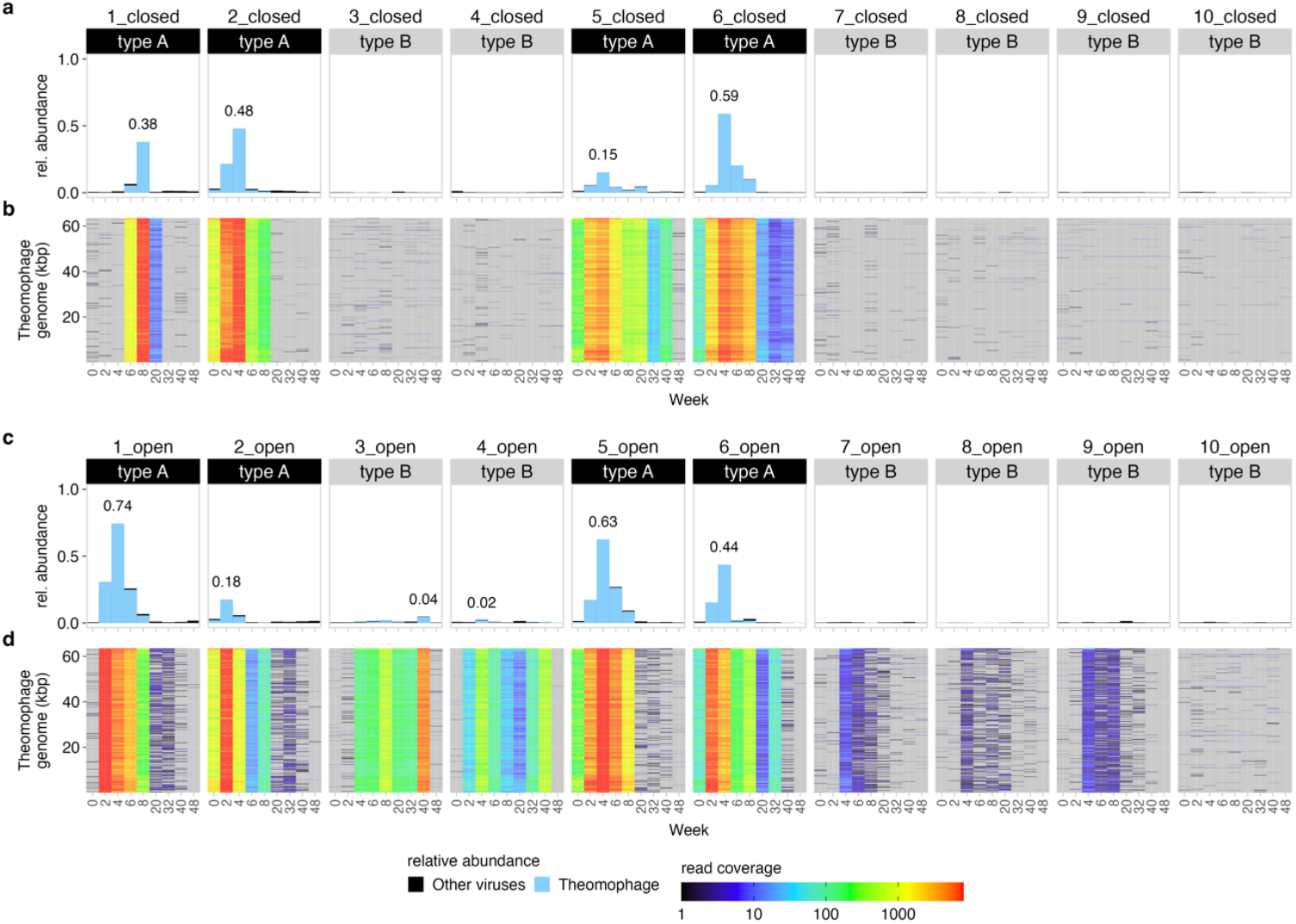
Massive outbreaks of a Theomophage, a novel Schitoviridae bacteriophage, are linked to community background. **a** Relative abundance of all predicted viral sequences (black) and Theomophage (blue), a highly abundant, previously undescribed bacteriophage belonging to the Schiroviridae family. Theomophage accounted for up to 59,1% and 74,3% of total community reads in closed (a) and open (c) samples, respectively. Read fractions >1% are labeled. **b** Genome coverage profiles show the percentage of sample nucleotides mapped to each position of the Theomophage genome. Theomophage was natively present in all closed type-A communities (1_closed, 2_closed, 5_closed, 6_closed), but absent or very rare in all closed type-B mesocosms. Grey indicates no mapped reads. **c** same as (b) for 10 open mesocosms. **d** same as (b) but for open mesocosms. Theomophage was detected in 5 out of 6 type-B mesocosms in the open regime, where it was absent in the corresponding paired closed mesocosms, indicating successful invasion in the open regime.

To determine whether Theomophage was present in type-B communities, we analyzed its genome coverage across all samples (**Fig.2b**; Methods). In closed type-B mesocosms Theomophage was absent or very rare, with no or very low genome coverage (type A: 69.2% ± 38.9%; type B 2.44% ± 2.18% genome breadth of coverage ≥1 read; mean ± SD; **Fig.S9, Table S2**). In contrast, Theomophage was detected in 5 of 6 open type-B communities, where it persisted until week 40 and reached up to 4.5% of total community reads (mesocosms closed_3, closed_4, closed_7-9, **Fig.2d**). This indicates that Theomophage, originally present in type-A communities, successfully colonized open type-B mesocosms via migration. Supporting this, the complete Theomophage genome was assembled from and detected in all sequenced MGE cocktail samples (**Fig.S10**). Despite successful colonization, Theomophage abundances were an order of magnitude lower in type-B communities than in its the native type-A context, indicating a strong effect of community type on ecological success. This observed lower abundance could be due to several factors. Host availability may have been lower in type-B mesocosms, or hosts may have exhibited higher levels of resistance. It is also possible that the original host of Theomophage was absent from type B, forcing it to infect a new, sub-optimal host. Although unlikely, due to the compositional nature of metagenomics, higher abundances of other taxa in type-B mesocosms could have reduced the relative abundance estimates of Theomophage, despite similar absolute abundances.

### Theomophage represents a new *Schitoviridae* subfamily

To determine the taxonomic affiliation of Theomophage, we constructed a gene sharing network using vContTACT2^39^ and the INPHARED viral reference set containing 26,302 bacterial and archaeal viral genomes (**Fig.3b**; Methods). Based on shared gene content, Theomophage grouped with *Schitoviridae* (formerly known as N4-like phages), a family of double-stranded DNA bacteriophages with podovirus-like morphology that are found globally in various habitats^40^. The Theomophage genome is organized in two large blocks on opposing strands, as is typical for *Schitoviridae*, and has 88 predicted genes and a single tRNA, including all seven proposed *Schitoviridae* hallmark genes^41^ (**Fig.3a**; **Table S3**, based on Pharokka^42^ and Phold^43^; Methods). Individual gene phylogenies of TerL, MCP, and portal protein also place Theomophage among the *Schitoviridae*, further supporting this taxonomic assignment (**Fig.S11-13**).

**Figure 3.**
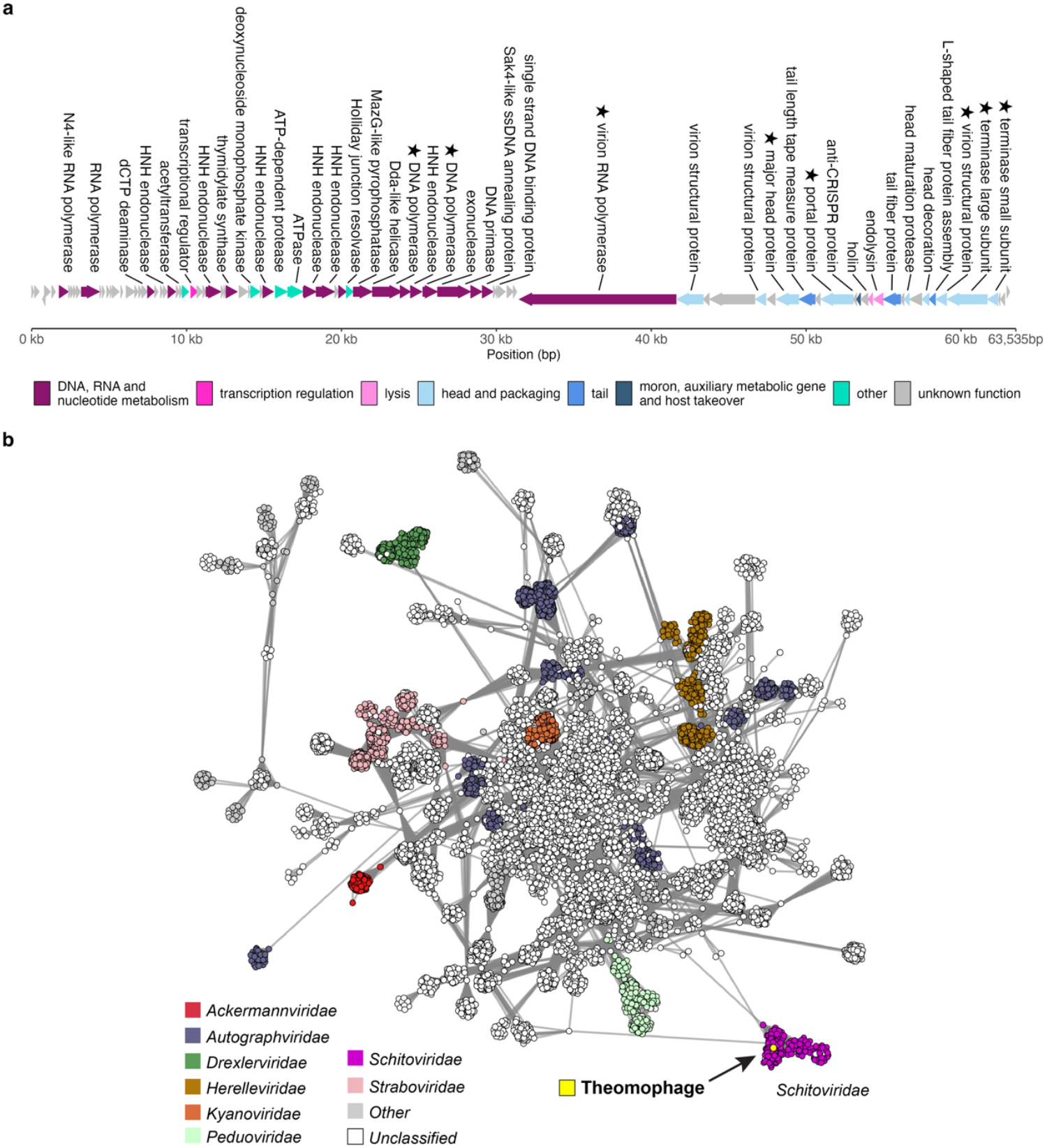
Genomic architecture and gene-sharing network of Theomophage, a novel Schitoviridae bacteriophage. **a** Theomophage genome (63,535 bp), encoding 88 genes and one tRNA, including all 7 Schitoviridae hallmark genes (indicated with stars). Gene functions for 40 genes were predicted with Phold^43^ and hallmark genes identified with HMM searches. **b** vConTACT2^39^ gene-sharing network of Theomophage and 26,302 bacterial and archaeal viruses from the INPHARED database^48^. Circles represent viral genomes, with edges indicating degree of connectivity based on shared protein clusters. Theomophage groups with bacteriophages of the Schitoviridae family (purple) but is only distantly related to known Schitoviridae sequences, suggesting it represents a new subfamily (**Fig.S14-S15**). Only the subnetwork containing Theomophage is shown.

To assess the phylogenetic placement of Theomophage within *Schitoviridae*, we analyzed 573 high-quality (>90% complete) *Schitoviridae* genomes and MAGs from three recent studies^40,41,44^. Using VIRIDIC^45^ to calculate intergenomic similarity (Methods), we found that the closest relative of Theomophage is IMGVR_UViG_3300045988_071216, a 72 kbp MAG co-assembled from human fecal samples, with only 8.98% intergenomic similarity (**Fig.S14**). This falls below the ≥20% threshold proposed for *Schitoviridae* subfamilies^41^, indicating that Theomophage represents a new subfamily. Gene-sharing analysis further supported this classification. Using vConTACT2, we found that Theomophage did not group with any of the 573 *Schitoviridae* genomes in vConTACT2 subclusters, which align with genus-level taxonomy as defined by the International Committee on Taxonomy of Viruses^39^ (**Fig.S15**; Methods). These results show that Theomophage is divergent from previously characterized *Schitoviridae* and represents a novel subfamily. To predict Theomophage lifestyle we used BACPHLIP^46^, which classified it as lytic (score 0.9625), which is common for Schitoviridae^41^. This is consistent with CheckV, which reported no bacterial flanking regions for any assembled Theomophage contig.

Cellulase activity has been reported for *Schitovirus* vB_EamP-S6 virions^47^, prompting us to investigate whether the high abundance of Theomophage in the cellulose-degrading communities might be related to cellulolytic capability. We queried the Theomophage genome using tblastn with vB_EamP-S6 cellulase genes Gp94, Gp95, and Gp96. No significant matches were found (**Table S11**), suggesting that the ecological success of Theomophage is not linked to cellulose degradation.

### Community composition remains stable despite massive Theomophage outbreaks

Viral predation can regulate microbial populations and drive major shifts in community composition and function^28,49^. For example, viral outbreaks following induced coccolithophore blooms can shift the balance between eukaryotic and bacterial organic matter recyclers^49^, and bacteriophage outbreaks are implicated in restoring gut microbial diversity following antibiotic treatment^50^. We hypothesized that the massive Theomophage outbreaks in type-A mesocosms would similarly impact community composition. However, Bray-Curtis dissimilarity between samples taken before and after Theomophage outbreaks was no greater than shifts over similar intervals in mesocosms without outbreaks (**Fig.S16**), suggesting that the communities were resilient to the Theomophage outbreaks. This observation may support the emerging view of a potentially limited impact of phages on bacterial community structure^2^.

### Candidate hosts of Theomophage

The high abundance of Theomophage in several compost mesocosms suggests that it likely infects a dominant community member. To identify potential hosts, we first searched for CRISPR spacers matching the Theomophage genome in all the compost metagenomes using Spacerextractor^51^ (Methods), recovering 298,447 spacers across mesocosm and cocktail datasets. None matched the Theomophage genome (up to 3 mismatches). A subsequent search against the external SpacerDB^52^ database revealed six spacers with 2–3 mismatches from *Nitrosomonas* MAGs (**Table S10**). While the high number of mismatches lowers confidence of a phage-host relationship, *Nitrosomonas* was abundant and strongly associated with type-A communities where Theomophage outbreaks occurred, and was also detected in 2 out of 5 open mesocosms that were invaded by Theomophage (open_3 and open_4; **Fig.1b–e; Fig.S17a**). Notably, Theomophage reached higher abundance in these two mesocosms than in the other three, providing further ecological support for a *Nitrosomonas* host. Although no direct spacer evidence (*i*.*e*. matching *Nitrosomonas* spacers) was found within the compost metagenomes, the spatial co-occurrence and spacer matches together suggest a possible host-phage association.

Two additional candidate hosts – *Moraxella*, and JACPRH01, a family within the *Nitrososphaerales* order of soil ammonia-oxidizing archaea^53^ – were identified through computational predictions. A single spacer with 1 mismatch was found in SpacerDB belonging to a *Moraxella* genome from a wastewater sample, while JACPRH01 was the highest-confidence prediction of iPHoP, a machine-learning framework that integrates multiple phage-host prediction tools (iPHoP score 85.80, corresponding to a 14.2% false discovery rate)^54–58^. Both taxa were detected in the compost metagenomes, but at very low abundance near the metagenomics detection threshold (a maximum of 182 reads or 0.0005**%** per sample) and showed no association with Theomophage presence (**Fig.S17bc**). If JACPRH01 is the true host, it would suggest an unexpected host range expansion of the *Schitoviridae* family to the archaeal domain, as all known *Schitoviridae* phages infect *Proteobacteria*^40^. Given the limited evidence and low abundance of these taxa, our best prediction remains a *Nitrosomonas* host, although further validation is required.

### Migration drives Theomophage evolution by exposure to new community contexts

The strong association of Theomophage with type-A communities, along with its successful colonization of type-B mesocosms, enabled us to investigate how migration and community context shape bacteriophage evolution. Specifically, we examined three conditions: (i) closed type-A mesocosms (no viral migration, native context), (ii) open type-A mesocosms (viral migration, native context), and (iii) open type-B mesocosms (viral migration, new context). We hypothesized that each condition would impose distinct selection pressures and result in different evolutionary signatures. In particular, we hypothesized that following colonization of non-native type-B communities, molecular evolution of Theomophage would be accelerated because its lower relative abundance suggests local maladaptation or a possible host switch. To test this, we identified Single Nucleotide Polymorphisms (SNPs) in Theomophage populations by mapping sample reads to reference assemblies (Methods). In total, we identified 76 SNPs with frequencies >10% in Theomophage populations. Open type-B mesocosms contained the greatest number of SNPs (n=70), and notably, 80.3% (n=61) of these variants were unique to type-B communities (**Fig.4ab, Fig.S18a**). This pattern supports the idea that exposure to new community contexts accelerates bacteriophage evolution and diversification.

**Figure 4.**
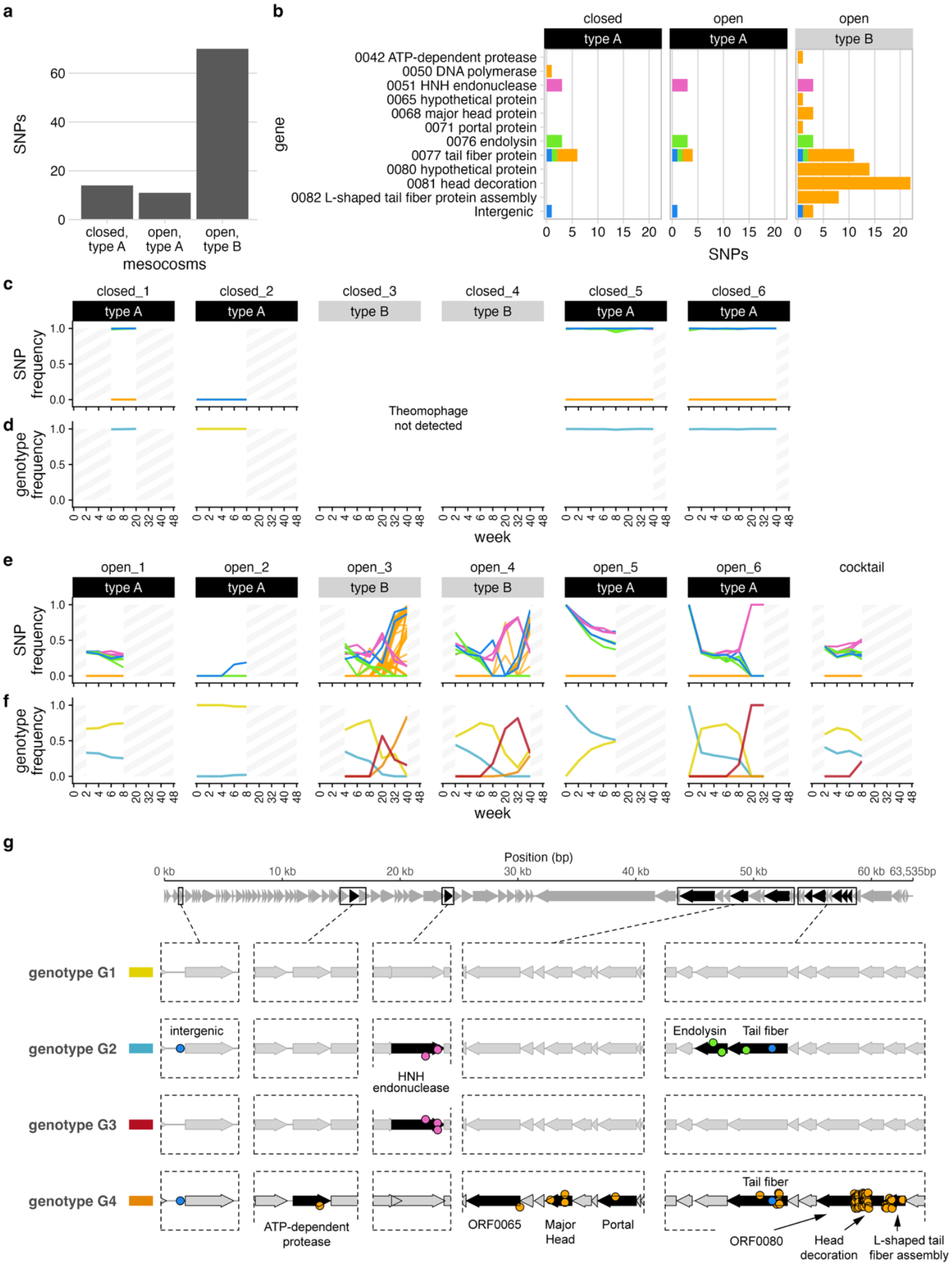
(next page) Molecular evolution and migration of Theomophage in closed and open mesocosms. **a** Total SNPs detected in Theomophage populations in type-A and type-B mesocosms under closed and open regimes. Most SNPs (80.3%) appeared only in the novel type-B communities, which Theomophage colonized in the open regime. **b** Genomic location of SNPs in (A). **c** Allele frequency trajectories of SNPs that differed from the reference genotype G1 in closed mesocosms. Theomophage populations in type-A communities were initially monomorphic for all variant positions, meaning they were composed of a single genotype: G1 in community closed_2, which was identical to reference, and G2 in communities closed_1, closed_5, and closed_6, which differed from the reference by 9 SNPs as indicated by frequencies of ~1. No new variants emerged that fixed in the population during the 40 weeks over which the bacteriophage was detected. Shading indicates samples with insufficient coverage for variant calling or where Theomophage was not detected. **d** Inferred genotype dynamics from (C); each mesocosm contained either G1 or G2. **e** Same as (C), but for open mesocosms and MGE cocktail. Both genotypes spread via the MGE cocktail, cross-invading type-A and previously Theomophage-free type-b mesocosms. Decoupled SNP trajectories at week 6-8 in mesocosms open_3 and open_6 indicate two independent recombination events between G1 and G2, giving rise to new genotypes G3 and G4. G4 also acquired 61 additional SNPs (orange). **f** Genotype dynamics in open mesocosms. Samples from open_7, open_8 and open_9 not shown due to insufficient read coverage for variant calling, although Theomophage was sparsely detected there (Fig.2). **g** Genomic positions of SNPs distinguishing genotypes G1, G2, G3 and G4. Most (46/61) new mutations in G4 occurred in ORFs 80, head decoration protein and L-shaped tail fiber assembly protein (**Table 1, Table S5**). SNPs are jittered vertically for clarity. SNP colors correspond to Fig.4b and Table 1 and are based on similarity of allele trajectories (Methods).

### Theomophage evolution is constrained in closed, native communities

Next, to explore how these SNPs emerged and spread in more detail, we reconstructed Theomophage genotypes and tracked them over time and across mesocosms. Reconstructing genotypes from short-read data is challenging because SNPS typically lie on different reads, making it hard to resolve linkage. Here we leveraged the observation that many samples were monomorphic at all variant positions, indicating clonal populations (i.e. all allele frequencies near 0 or 1; Methods). Using this approach, we identified two genetically distinct Theomophage genotypes in the founding communities. Mesocosm 2 contained a genotype identical to the assembled vMAG used for variant calling (hereafter genotype G1), while mesocosms 1, 5, and 6 harbored a second genotype (G2) which differed from G1 by nine SNPs located in genes associated with host interaction and replication, including the tail fiber protein, HNH endonuclease, endolysin, and one intergenic region (**Fig.4c,g**; **Table 1**). Each founding mesocosm contained a single genotype, with minimal within-population diversity.

**Table 1.** Genomic variants defining four Theomophage genotypes. For each Single Nucleotide Polymorphism (SNPs), the position, effect, associated open reading frame (ORF), predicted gene function, and inferred genotype for each genotype are shown. Nine SNPs differentiate genotype G2 from G1. Details for SNPs arising in G4 are provided in Supplementary Table 5. Colors match Fig.4b-g.

| Variant | Effect | ORF | Predicted Function | G1 | G2 | G3 | G4 | Color in Fig.4 |
| --- | --- | --- | --- | --- | --- | --- | --- | --- |
| 1276 T → G | intergenic | – | – | – | ✓ | – | ✓ | — |
| 24224 T → C | Ile → Thr | 0051 | HNH endonuclease | – | ✓ | ✓ | – | — |
| 24375 G → A | synonymous | 0051 | HNH endonuclease | – | ✓ | ✓ | – | — |
| 24378 T → C | synonymous | 0051 | HNH endonuclease | – | ✓ | ✓ | – | — |
| 54689 T → C | Glu → Gly | 0076 | endolysin | – | ✓ | – | – | — |
| 54853 T → C | synonymous | 0076 | endolysin | – | ✓ | – | – | — |
| 54856 C → T | synonymous | 0076 | endolysin | – | ✓ | – | – | — |
| 55320 A → C | Val → Gly | 0077 | tail fiber protein | – | ✓ | – | – | — |
| 55814 C → T | Synonymous | 0077 | tail fiber protein | – | ✓ | – | ✓ | — |
| 16296 T → A | Leu → Ile | 0042 | ATP-dependent protease | – | – | – | ✓ | — |
| 22232 A → G | Synonymous | 0050 | DNA polymerase | – | – | – | ✓ | — |
| 46679 C → T | Gly → Ser | 0065 | hypothetical protein | – | – | – | ✓ | — |
| 3 SNPs | see Table S5 | 0068 | major head protein | – | – | – | ✓ | — |
| 51895 G → T | Leu → Ile | 0071 | portal protein | – | – | – | ✓ | — |
| 14 SNPs | see Table S5 | 0077 | tail fiber protein | – | – | – | ✓ | — |
| 14 SNPs | see Table S5 | 0080 | hypothetical protein | – | – | – | ✓ | — |
| 2 SNPs | intergenic | – | – | – | – | – | ✓ | — |
| 22 SNPs | see Table S5 | 0081 | head decoration | – | – | – | ✓ | — |
| 8 SNPs | see Table S5 | 0082 | L-shaped tail fiber protein assembly | – | – | – | ✓ | — |

With the identity and initial distribution of genotypes established, we next tracked how these genotypes changed over time within each mesocosm. In closed mesocosms, no new SNPs appeared in either genotype G2 or G1 over the 40 weeks in which Theomophage was detected, though genotype G1 contained limited preexisting microdiversity in mesocosms closed_5 and closed_6 (**Fig.4c, S18**). The absence of novel mutations is striking given the massive phage outbreaks and the long duration over which Theomophage was observed, which suggest extensive viral replication. In experimental evolution studies in simplified phage–host systems^4,19^ and dairy starter cultures with limited taxonomic diversity^59^, phage evolution is typically rapid and driven by coevolution. In contrast, we observed prolonged genetic stability under ecologically complex conditions. Consistent with previous observations in natural environments^14,15,60^, our results suggest that phage evolution in complex microbial communities may proceed only slowly, with long-term, stable coexistence between phage and host.

### Migration accelerates evolution via recombination between preexisting Theomophage genotypes

In contrast to the genetic stability observed in closed mesocosms, Theomophage populations in open mesocosms exhibited rapid and complex evolutionary dynamics, driven by genotype migration, recombination, and the acquisition of new SNPs following colonization of the type-B mesocosms. While closed mesocosms only contained either genotype G1 or G2, in open mesocosms both genotypes were often detected immediately following the first serial transfer, indicating rapid dissemination between mesocosms. (**Fig.4ef**). For example, mesocosm open_5 initially contained only genotype G2, but by week 2 genotype G1 had invaded and rose to ~50% frequency by week 8, as indicated by the coupled frequency decline of the nine SNPs that characterized G2. Similar cross-invasions were observed in all other open mesocosms with sufficient coverage for variant calling, including colonization of the novel type-B mesocosms by both Theomophage genotypes. Importantly, both migrating genotypes were also detected in the MGE cocktail used for transfers between mesocosms, providing further evidence that these patterns resulted from viral migration rather than parallel evolution (**Fig.4ef**).

In all open mesocosms where G1 and G2 coexisted beyond week 8, the frequency trajectories of their characteristic alleles became decoupled, indicating recombination between the genotypes. For example, in open_6, genotype G1 and G2 coexisted for 8 weeks, after which a recombinant genotype (G3) emerged, characterized by three G1-derived alleles in HNH endonuclease and six G2-derived alleles (**Fig.4ef**; Table 1). Similarly, in open_3 a second recombinant genotype (G4) arose that combined alleles from both founder genotypes. Both recombinants replaced their parental genotypes and spread to other mesocosms via continued migration. G3, originating in open_6, was detected in open_3 and open_4, while G4 spread from open_3 to open_4 and eventually to open_6 by week 32, as revealed by a more sensitive SNP analysis (**Fig.S19**).

### Rapid SNP accumulation following migration to new community type

At week 8, 61 new SNPs appeared in open_3 that were not detected in any founding or closed communities (**Fig.4ef**). The close similarity in frequency trajectories between these novel SNPs and the defining alleles of G4 indicates that they were located on the same genome. These mutations occurred in several predicted structural proteins, including Major Capsid and Tail proteins, but were mostly clustered (46/61) in a region spanning three consecutive genes: ORF0080, head decoration protein, and L-shaped tail fiber assembly protein (**Fig.4bg, Table S5**).

To investigate the mechanistic origin of these mutations, we first tested whether these mutations arose *de novo* via diversity-generating retroelements (DGRs)—error-prone reverse transcriptase systems that target specific genomic loci, typically structural genes, to promote rapid adaptation to new hosts^61^. However, screening the Theomophage genome with MyDGR^62^ revealed no DGRs or reverse transcriptase genes, making this mechanism unlikely. Another major driver of viral evolution is recombination^12^. The tight genomic clustering of mutations suggests that recombination, rather than independent point mutations, may underlie the emergence of these variants. However, a blastn^63^ query of all assembled contigs with the Theomophage region spanning the three mutated genes yielded no significant matches. This suggests that recombination may have occurred with a rare sequence not captured in our assemblies. Rare viral genotypes (e.g. below the metagenomics detection threshold) can act as viral seed banks in natural ecosystems and harbor genetic diversity that can become adaptive when selection pressures change^64^. We hypothesize that following migration to type-B mesocosm open_3, Theomophage recombined with a member of the rare virosphere giving rise to genotype G4. While the precise mechanism remains unresolved, the timing, localization, and spread of these mutations from open_3 to open_4 and open_6 suggests they conferred selective advantage in the type-B community context.

## Discussion

The factors that drive bacteriophage evolution in complex natural ecosystems, such as local microdiversity, community context, and migration, cannot be fully captured by laboratory studies that reduce the genomic and spatiotemporal complexity of natural environments^10,16^. Previous studies have examined the effect of migration using single phage-host pairs^65,66^, but its influence in taxonomically rich communities has not been explored.

Our results show that bacteriophage population dynamics and evolution in complex communities are highly contingent on community background and migration. In isolated, native mesocosms, Theomophage showed no detectable molecular evolution over 40 weeks, despite repeated massive outbreaks. In contrast, migration to new communities sparked rapid evolution via recombination between preexisting genotypes and SNP accumulation, producing new genotypes that spread across mesocosms.

Our results indicate that in complex microbial communities, phage–host dynamics can remain evolutionarily static for 40 weeks unless disrupted by migration. This contrasts with results from simplified lab systems, where antagonistic coevolution often produces rapid phage evolution via sequential selective sweeps^8–10^. Instead, our findings match diversity patterns in natural microbial ecosystems that indicate long-term coexistence of susceptible and resistant hosts without rapid turnover^14,15^. They further suggest that in natural ecosystems, disturbances to local community structure such as seasonal rainfall in soils^18,67^ or cycles of dispersal and community assembly on marine organic particles^68^, may act as major drivers of bacteriophage diversification.

Viral abundance is a key parameter in microbial ecology but remains challenging to measure in complex microbial communities^69^. Two methods are commonly used: microscopy-based counting of virus-like particles (VLPs), which estimates free viral particles but cannot detect ecologically relevant intracellular viral stages or provide taxonomic or functional insight; and metagenomics, which captures both free and intracellular viruses and enables genome-level characterization. Each method has limitations^69,70^, but metagenomics is uniquely suited to quantify specific viral taxa in community context^71^. Here, using metagenomics, we have captured evidence of massive, parallel outbreaks of Theomophage, a previously undescribed bacteriophage, across different independent compost communities, where it reached unprecedented abundances of up to 74.3% of metagenome reads. Such dominance by a single virus is rare: viral reads typically constitute ~5% of metagenomic sequences^72^, though higher values have been reported. For example, crAssphage can reach 22% in human gut metagenomes^73^ and a *Streptococcus* phage comprised ~20% of reads in cheese starter cultures composed of two bacteria and three associated phages^74^. Our findings far exceed these cases, showing that a single phage can dominate even highly diverse microbial communities in terms of genomic information.

The extraordinary abundance of Theomophage highlights the need to account for viral sequences when analyzing metagenomes. State-of-the-art taxonomic profilers used in metagenome analysis struggle to identify viral sequences^75^, requiring the use of specialized viral identification tools^76,77^. Comprehensive viral profiling remains limited to well-described biomes^72^, meaning that the most abundant members of microbial communities may be missed in metagenome profiling.

Viral abundance estimates using metagenomics can be influenced by technical biases during DNA extraction, library preparation, and sequencing^69,70,78^. In particular, rolling circle amplification is known to skew viral abundances^79^, but this was not used here. The Nextera XT library preparation kit used in this study may underrepresent low-GC regions^80^, but given that Theomophage has a lower GC content (34%) than the average of the assembled metagenomes, this bias would underestimate rather than inflate its abundance. Thus, our Theomophage abundance estimates may be conservative.

One potential explanation for the exceptional abundance of Theomophage is a carrier state-like infection mode, in which cells shed virions without undergoing lysis. This non-canonical infection mode has been observed in crAss-like phage crAss001 where it was proposed to underlie high abundance and stable coexistence with a *Bacteroides* host^81^. Similar modes have been reported for a wide range of virulent phages^82–87^, and it is possible that a comparable mechanism enabled the high abundances and long-term persistence of Theomophage reported here.

Finally, we highlight potential limitations of our study. First, we cannot rule out that the coarse temporal resolution in later stages of the experiment caused us to miss additional Theomophage outbreaks. The more frequent sequencing earlier in the experiment showed that outbreaks took 6 to 8 weeks, suggesting similar events may have occurred undetected later on. The temporal resolution also limited our ability to assess the impact of Theomophage outbreaks on community composition. Second, a confirmed host would provide valuable context for interpreting the evolutionary dynamics of Theomophage. Host identification could clarify whether the observed genotype-level diversity in native type-A communities reflects local adaptation, and whether the increased evolutionary rates following migration to new community types are mirrored in host evolution, or a signature of adaptation to a new, suboptimal host^88^. If *Nitrosomonas* were confirmed as the host, this could suggest a host switch following migration to mesocosms open_7, open_8 and open_9, where *Nitrosomonas* was sparsely detected.

Together, our study has leveraged genotype-resolved analysis of existing shotgun metagenomic data to reach new conclusions about the ecology and evolution of a previously unknown bacteriophage in a complex microbial ecosystem. We identified two alternative, stable community types, observed massive and reproducible bacteriophage outbreaks confined to one type, and showed that phage migration and colonization of the second community type accelerated bacteriophage evolution. These findings demonstrate that genotype tracking of laboratory-propagated natural microbial communities can reveal viral eco-evolutionary dynamics with genomic precision, providing a powerful approach for studying viruses in complex yet tractable ecosystems.

## Materials and Methods

Note that the closed and open treatments were previously referred to as “vertical” and “horizontal”, respectively^30,34^.

### Assembly and binning

Read quality control was done with Prinseq v.0.20.4^89^ using parameters -derep 14 -lc method dus -c threshold 20. Adapters were trimmed using Flexbar v.3.5.0^90^ with -- adapter-preset Nextera -ap ON. The high complexity of compost ecosystems makes metagenome assembly difficult, potentially leading to short contigs and a large fraction of unassembled reads^91^. To improve assembly and increase the chance of recovering low-abundant genomes such as bacteriophages^92,93^, reads from all timepoints for each mesocosm were pooled and co-assembled with metaSPAdes v.3.14.0^94^. To preserve sample-level diversity for Theomophage genome annotation and variant calling, all samples (n=170) were also assembled separately. Samples from the viral cocktails were assembled in the same way. Reads were mapped using BWA-mem2 v0.7.17-r1188^95^ and coverage was calculated with SAMtools v1.7^96^. Contig depths were calculated with jgi_summarize_bam_contig_depths and binned using Metabat2 v.2.12.1^97^.

### Taxonomic profiling

Taxonomic classification of assembled scaffolds and bins was performed with CAT v5.3^31^ with --index_chunk 1 --block_size 5 --top 11 --I_know_what_Im_doing, using CAT NCBI database and taxonomy v2021-04-30. Taxon abundances (fraction of total sample reads) were calculated using RAT version Sept 2021^32^ and database v2021-04-30. For dimension reduction, unclassified taxa or taxa that were only classified at other taxonomic ranks were dropped, and relative abundances were center-log-ratio (CLR) transformed, with zero values adjusted by adding 1 to all read counts. To estimate community richness the number of detected genera was counted with different abundance cutoffs (**Table S1**). Reads from contigs that could not be annotated at genus level by RAT, for example due to conflicting information from ORFs or incomplete taxonomic annotation of best matches, were either counted as a separate genus, or removed. Putative viral sequences were identified with WhatThePhage v0.9.0. While counting any contig marked as viral by this panel of virus identification tools likely increased the number of false positives, our subsequent analysis focused on a single bacteriophage sequence identified with high confidence by manual curation. To estimate viral abundances (including Theomophage), we randomly selected a co-assembly where Theomophage was completely assembled and used that as a target for read mapping.

### Viral completeness estimation, annotation, lifestyle prediction

Viral contig completeness was estimated using CheckV v1.0.3 with database v1.5^38^. Theomophage contig NODE_224 was estimated 100% complete based on the presence of 55bp direct terminal repeats (DTRs). Complete Theomophage genomes were independently assembled in 54 independent, single-timepoint assemblies, with a consistent contig length of 63,535bp (46/54 contigs, **Fig.S20)** and high sequence similarity (99.98 ANI). The genomes were circularly permuted and contained DTRs, consistent with complete genome assembly from shotgun reads derived from phage replication concatemers or linear genomes with DTRs (**Fig.S21**).

Theomophage MAG NODE_224_length_63535 assembled from sample 3_open-week4 was annotated with Pharokka v1.7.5^42^ and Phold v0.2.0^43^ using default settings. Specifically, coding sequences (CDs) were predicted with PHANOTATE^98^, tRNAs were predicted with tRNAscan-SE 2.0^99^, tmRNAs were predicted with Aragorn^100^ and CRISPRs were predicted with CRT^101^. Functional annotation was generated by translating protein amino acid sequences to the 3Di token alphabet with ProstT5^102^, and Foldseek^103^ was then used to search these against the Phold phage protein structures database, which contains PHROGs^104^, VFDB^105^ and CARD^106^ structures predicted with Colabfold^107^. To search for the seven *Schitoviridiae* hallmark genes, a protein hidden Markov model (HMM) was constructed for each from protein sequences from REF^41^ by aligning them with MAFFT v7.526^108^ and hmmbuild (HMMER v3.4, www.hmmer.org). The resulting models were used to search the predicted Theomophage CDs using hmmsearch (HMMER v3.4) with default parameters. Output is available as **Table S4. Fig.3a and 4g** were drawn in R v4.3.1 with gggenes v0.4.1 (https://github.com/wilkox/gggenes) and ggmagnify v0.4.1 (https://github.com/hughjonesd/ggmagnify). Theomophage lifestyle was predicted with BACPHLIP v.0.9.6^46^.

### Theomophage taxonomy

The gene sharing network was created by combining predicted proteins of 26,302 bacterial and archaeal viral reference genomes from the INPHARED reference set (1Oct2023)^48^ with predicted proteins from Theomophage NODE_224_length_63535, using vConTACT2 v.0.9.19 (–rel-mode ‘Diamond’ –pcs-mode MCL –vcs-mode ClusterONE). For **Fig.S15** a gene sharing network was created for Theomophage and 573 *Schitoviridae* reference sequences (see below). Networks were visualized with Cytoscape v.3.9.1^109^ and Adobe Illustrator 2025).

A reference set of 573 high quality (predicted >90% complete) *Schitoviridae* sequences was created by downloading sequences described as “high quality” or “complete” in Wittmann *et al*^40^ and Zheng et al^41^ from Genbank and IMG/VR v3, complemented by adding sequences annotated as “Schitoviridae” and “high quality” or “isolate” from IMG/VR v4^44^, finally followed by dereplication at 100% ANI with CD-HIT v.4.8.1^110^. For gene sharing networks, protein prediction was performed as described above.

For tailed bacteriophages three conserved proteins - the Terminase large subunit (terl), major capsid protein (MCP) and portal protein (portal) can be used to infer taxonomic relationships^111^. We gathered proteins annotated as “terminase large”, “portal”, or “major capsid” or “large capsid” from the INPHARED dataset and added the corresponding protein from Theomophage MAG NODE_244. Genes were aligned with Clustal Omega v.1.2.4^112^ using default parameters and positions with gaps in more than 50% of the sequences were removed with trimAl v.1.4.rev15^113^. Gene trees were made using FastTree version 2.1.10^114^ with default parameters and plotted with R v.4.3.1 using the ggtrees package^115^.

Intergenomic similarities (pairwise nucleotide identity normalized for genome lengths) were calculated from MAG NODE244 and 573 high-quality *Schitoviridae* sequences using VIRIDIC standalone v1.1 (blastn parameters “-word_size 7 -reward 2 -penalty -3 -gapopen 5 -gapextend -2”)^45^.

### Host prediction

iPHoP v1.3.3, database Aug_2023_pub_rw^54^ was run with default parameters and additionally with a lowered confidence cutoff (−m 80). iPHoP predictions and raw output per bioinformatic tool are available as **Table S6-S7**.

Crispr spacers were extracted from mesocosm and cocktail reads and filtered and denoised with Spacerextractor^51^ v.0.9.5 using default parameters. Resulting spacers were mapped against the four inferred Theomophage genotypes (MAG NODE_224_length_63535_cov_5745_934389, genotypes G2, G3 and G4) using Spacerextractor with default parameters, which resulted in no matches. Spacers from the global SpacerDB database v.2025-05-02^52^ were mapped in the same way against the four Theomophage genotypes, only considering spacers that mapped with 0 or 1 mismatch across the full spacer length. Results are available as **Table S8-S10**.

### Variant calling

To reduce the chance of spurious read mapping, QC-filtered reads were competitively mapped against all contigs from a randomly selected single-sample assembly containing the complete Theomophage MAG (sample open_3-week_4 containing NODE_224_length_63535_cov_5745_934389) with BWA-mem2^95^. Variants were called for the Theomophage MAG using Haplotypecaller, GenomicsDBImport and GenotypeGVCFs from GATK v.4.2.3.0^116^ following the joint variant calling pipeline and the best practices for germline short-variant discovery. To exclude sequencing and read mapping artifacts, only alleles that reached a frequency of 10% in any sample and were supported by a minimum of 4 reads, coming from samples where ≥30% of the genome was covered by ≥10 reads were considered for downstream analysis. This resulted in a total of 76 unique SNPs across all samples. To further exclude read mappings artifacts, read depth coverage profiles were inspected with Integrative Genomics Viewer V.2.13.2^117^. Variants located in genomic areas with irregular coverage (e.g. abrupt depth changes) were further evaluated by separately reassembling mapped read pairs for these samples and a second round of variant calling using the resulting contig as reference. All SNPs were confirmed and no variants were removed.

Variant effects were predicted with SNPEff v.5.0e^118^ (-noLog -noStats -no-downstream -no-upstream -no-utr) using a custom SNPEff database built with NODE_224_length_63535_cov_5745_934389 as reference, a .gff file generated using Prokka and the Bacterial_and_Plant_Plastid codon table. MAGs were interrogated for Diversity Generating Retroelements (DGRs) with the webserver MyDGR^62^.

### Resolving genotypes and estimating genotype abundance

Genotypes were inferred from samples where variant frequencies for all 76 variants were close to either 0 or 1, indicating monomorphic populations. For example, genotype G1 was inferred from sample closed_2 week 0, where the frequency of all 76 variants was close or equal to 0, indicating that this population consisted of a single genotype that was identical to the reference genome (**Fig.4c,d,g; Table 1**). Similarly, genotype G2 was inferred from monomorphic samples closed_1, closed_5, closed_6 week 0, which had nine SNPs in the tail fiber protein, HNH endonuclease, endolysin and an intergenic position with frequencies near 1. Note that Theomophage populations were not monomorphic in open samples, but the 9 SNPs that differentiated S1 and G1 showed similar frequency dynamics over time, further suggesting genetic linkage in these samples in accordance with the pigeonhole principle. Similar reasoning allowed the identity of G3 to be inferred from samples open_6 Week 20 and 32, and G4 from sample open_3 week 40. For **Fig 4.D** and **F** we then assumed the simplest scenario that could explain the evolution of G3 and G4 from G1 and G2 that was compatible with the variant trajectories, namely two separate recombination events and fixation of 61 new variants in G4. We approximated genotype abundances for weeks 8, 20, 32 and 40 in communities open_3, open_4, open_6 and the cocktail as follows: G4 abundance as the average relative abundance of orange and blue variants; G3 abundance as the average of pink variants; G2 as the average of the blue and green variants, and G1 as 1 minus the abundance of G2, G3 and G4. For other time points the abundance of G2 was estimated as the average of blue, pink, and green SNPs and G1 as 1 minus the abundance of G2. The four genotypes inferred above accounted for 92.1% of the reported SNPs and we conclude that they captured most of the relevant dynamics of genetic diversity within the Theomophage population. The remaining 6 SNPs represent further community-specific microdiversity within genotype G2 that was limited to communities closed_5, closed_6 and open_5 **(Fig.S21)**. For visual clarity we gave SNPs with similar trajectories from cluster 1 different colors.

Unless noted otherwise, figures and analysis was done in RStudio v.2022.12.0 with R v.4.2.0 using cowplot v1.1.1 (github.com/wilkelab/cowplot), Tidyverse v.1.3.2^119^, data.table v1.14.6, ggbeeswarm v.0.7.1.

## Author Contributions

JM wrote the paper with input from PH and BED. JM, PH, BED contributed equally to design of the analysis and interpretation of the results. JM performed the data analysis. PS wrote code used for variant calling and advised with the variant calling analysis. PBR provided the original experimental design and ensuing data and commented on the manuscript.

## Data availability

All data and code will be made publicly available before publication. Code for analysis and figure generation will be made available on github.com/MGXlab/theomophage. The Theomophage genome will be deposited in Genbank. Raw sequencing data and assemblies will be made available through online repositories.

## Acknowledgments

The authors gratefully acknowledge Jan Kees van Amerongen for maintaining the computational facilities. JM is supported by the Dutch Research Council (Nederlandse Organisatie voor Wetenschappelijk Onderzoek, www.nwo.nl) grant 022.005.023 and the Utrecht University BINF fund. BED is supported by European Research Council (ERC) Consolidator grant 865694: DiversiPHI, the Deutsche Forschungsgemeinschaft (DFG, German Research Foundation) under Germany’s Excellence Strategy – EXC 2051 – Project-ID 390713860, the Alexander von Humboldt Foundation in the context of an Alexander von Humboldt-Professorship founded by German Federal Ministry of Education and Research, and the European Union’s Horizon 2020 research and innovation program, under the Marie Skłodowska-Curie Actions Innovative Training Networks grant agreement no. 955974 (VIROINF). PBR acknowledges core support from the Max Planck Society and funding from the Deutsche Forschungsgemeinschaft (DFG) Collaborative Research Center 1182 ‘Origin and Function of Metaorganisms’ (grant no. SFB1182, Project C4).

