## Supplementary Material for "Eco-evolutionary dynamics of massive, parallel bacteriophage outbreaks in compost communities"

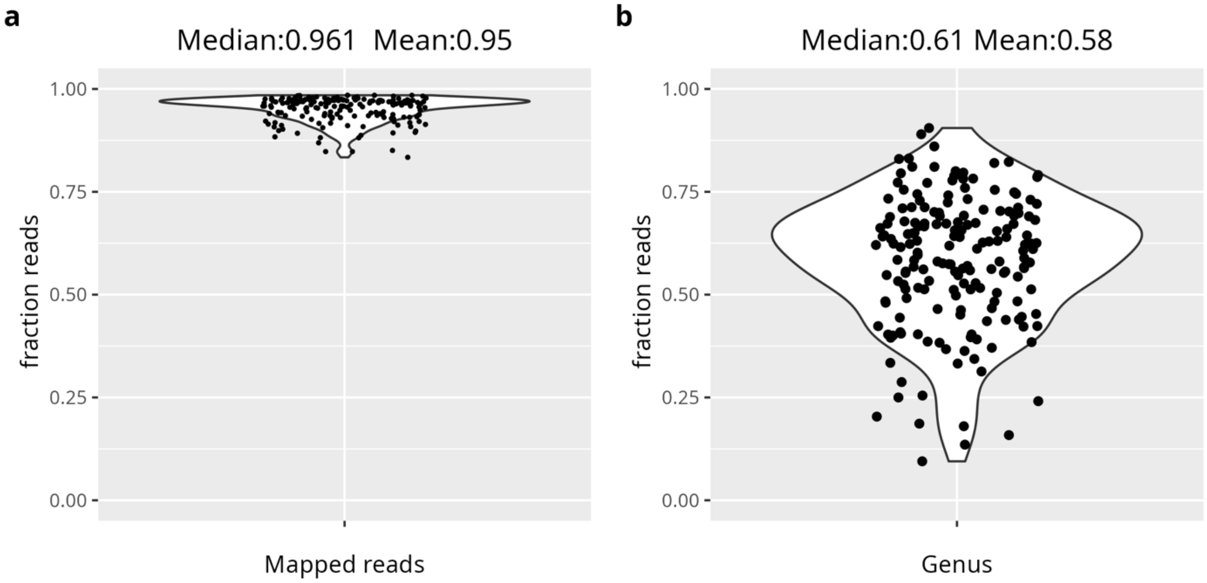


**Figure S1. a.** Fraction of sample reads mapped to contigs (170 samples). **b.** Fraction of mapped sample reads annotated at genus rank.

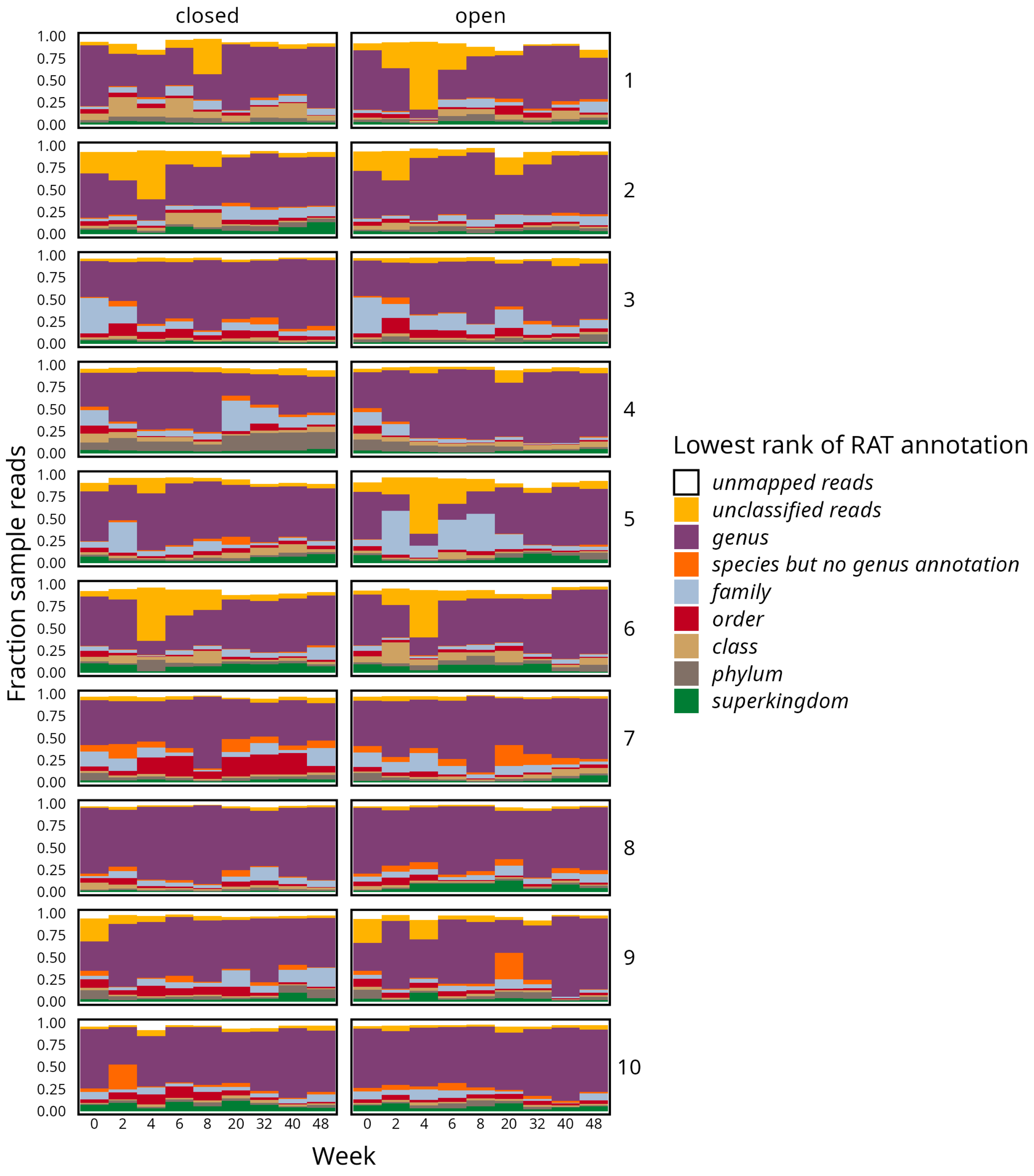


**Figure S2.** Fraction of sample reads annotated at genus rank, or alternatively the lowest available taxonomic rank. Reads in orange are annotated at species but not genus rank, due to the the top-scoring RAT match lacking genus annotation in the NCBI non-redundant protein database. Rows indicate mesocosms 1-10.

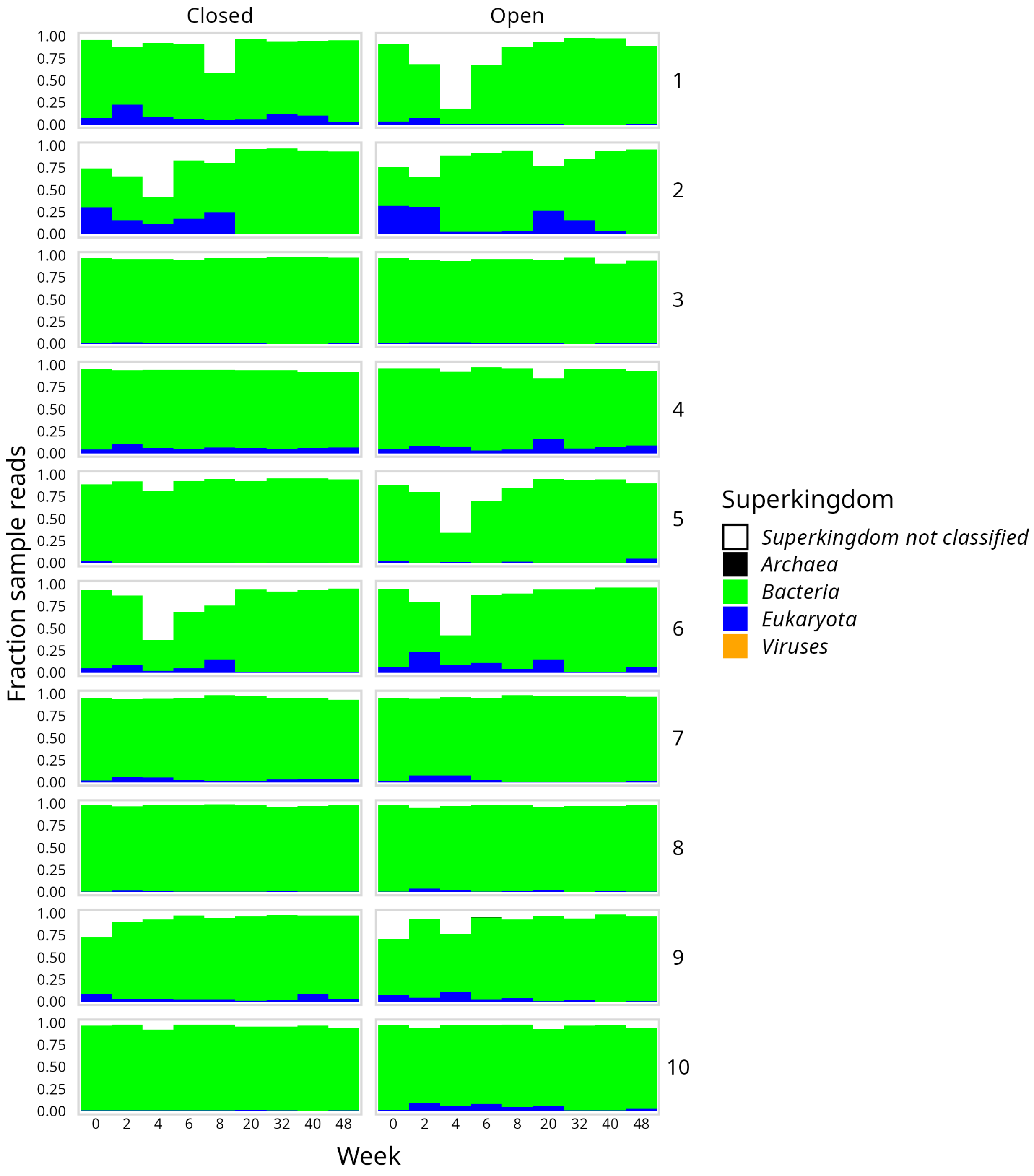


**Figure S3.** Fraction of sample reads annotated at superkingdom rank, annotated with RAT (see Methods for details). Rows indicate mesocosms 1-10.

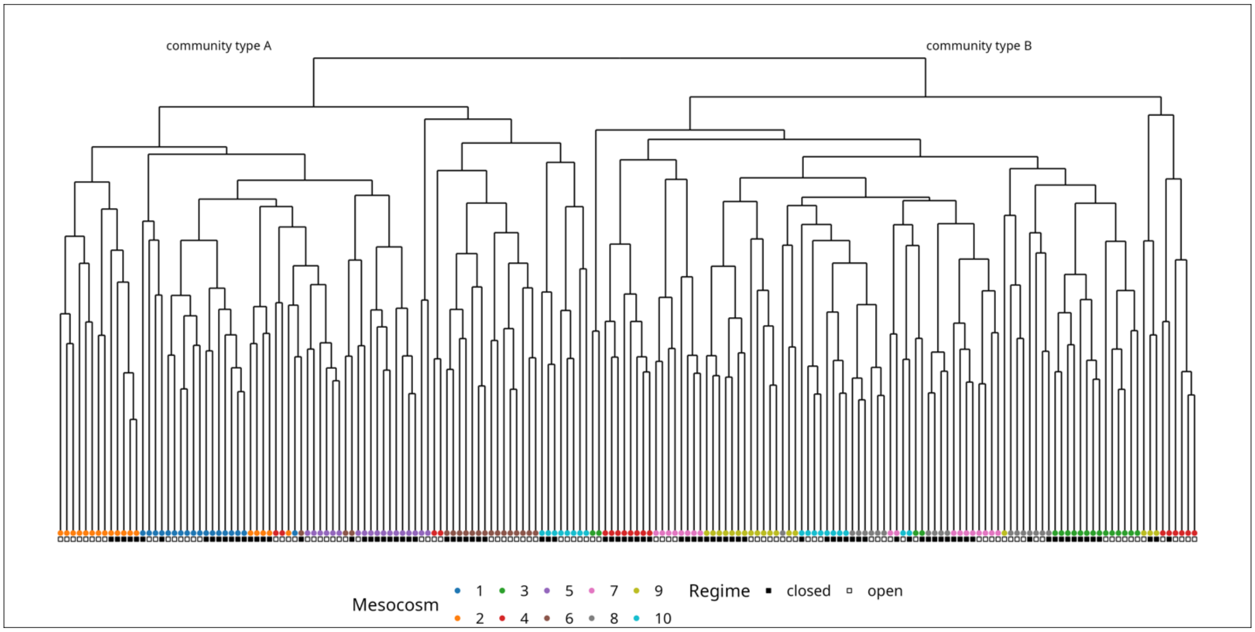


**Figure S4.** Hierarchical clustering of mesocosm samples based on Aitchison distance of genus-level abundances. Samples cluster into two distinct community types, consistent with the PCA shown in Fig. 1d of the main text (see Methods for details).

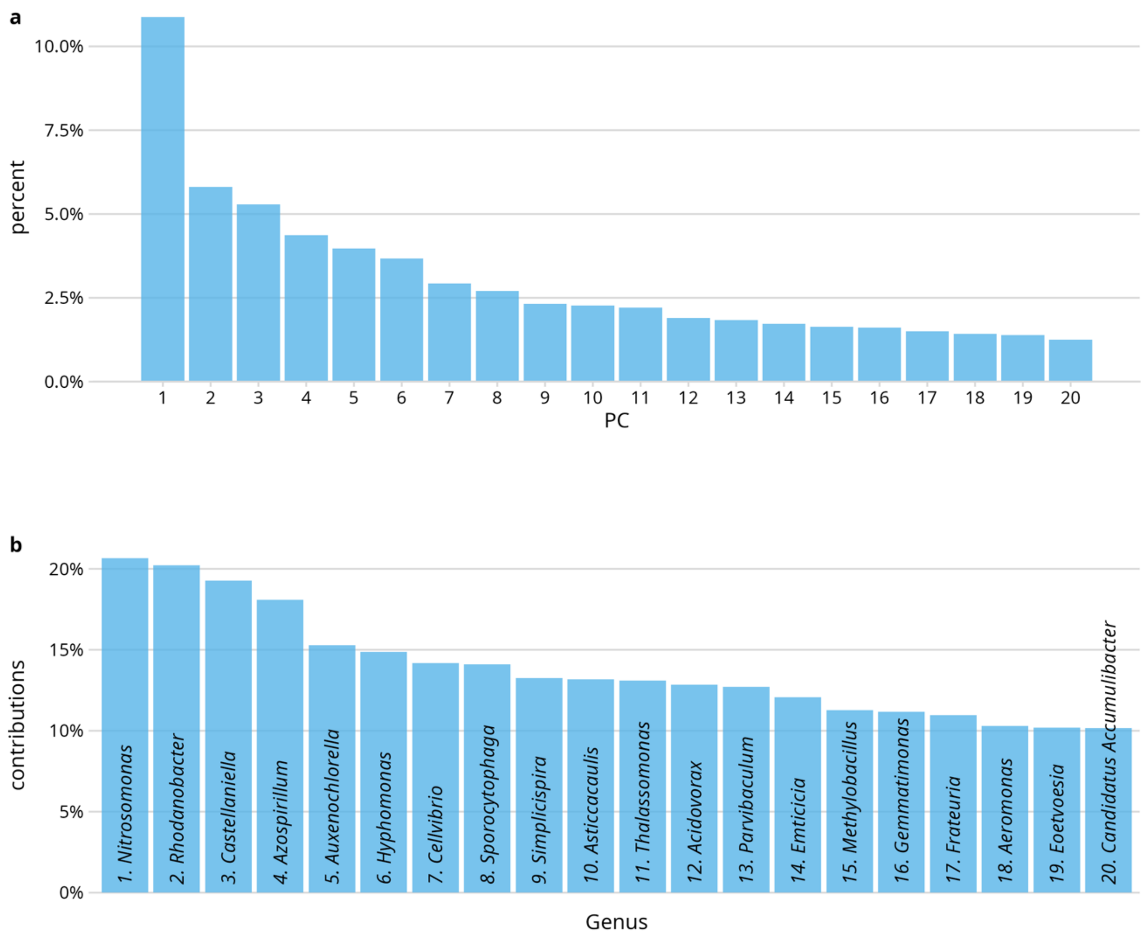


**Figure S5.a.** Variance explained by the top 20 principal components in the PCA shown in Fig. 1d (main text). **b.** Top 20 genera contributing to PC1 and PC2.

**a**

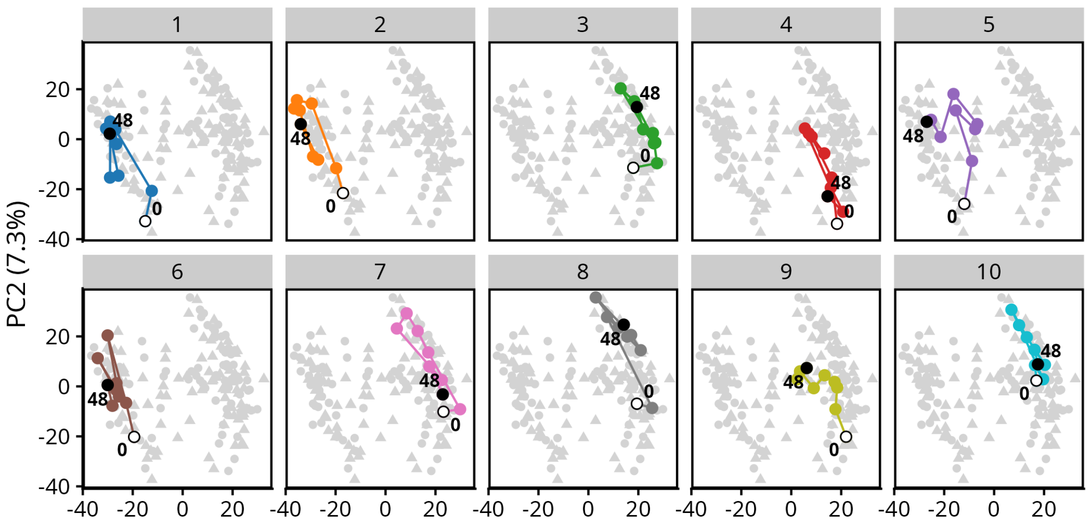


**b**

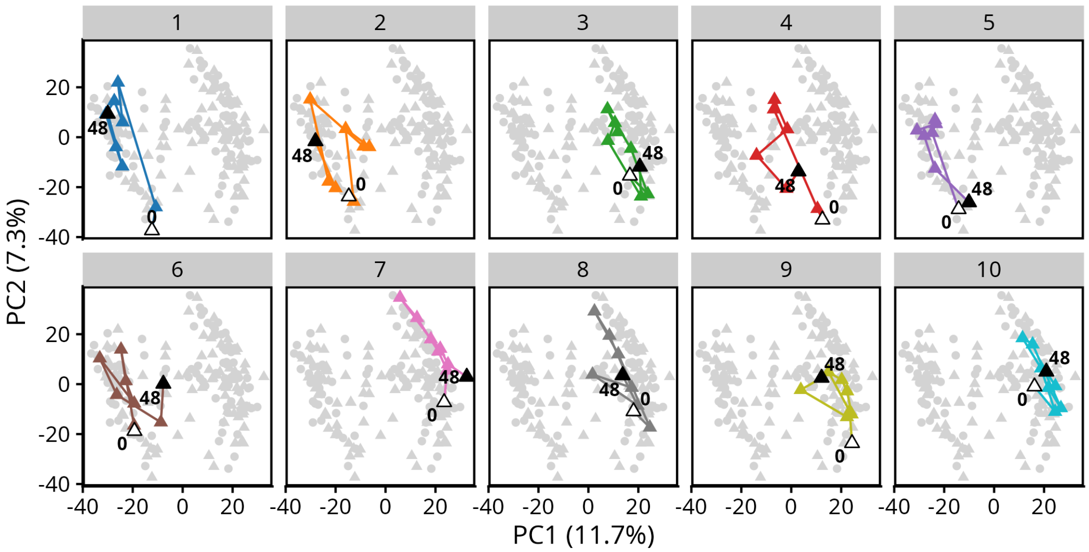


**Figure S6.a.** PCA of closed mesocosms (closed_1–closed_10), taxonomically profiled at genus rank. Consecutive samples are connected by lines per mesocosm. Open symbols indicate the starting point (week 0), and closed symbols indicate the final time point (week 48). **b.** Same as (A), but for open mesocosms.

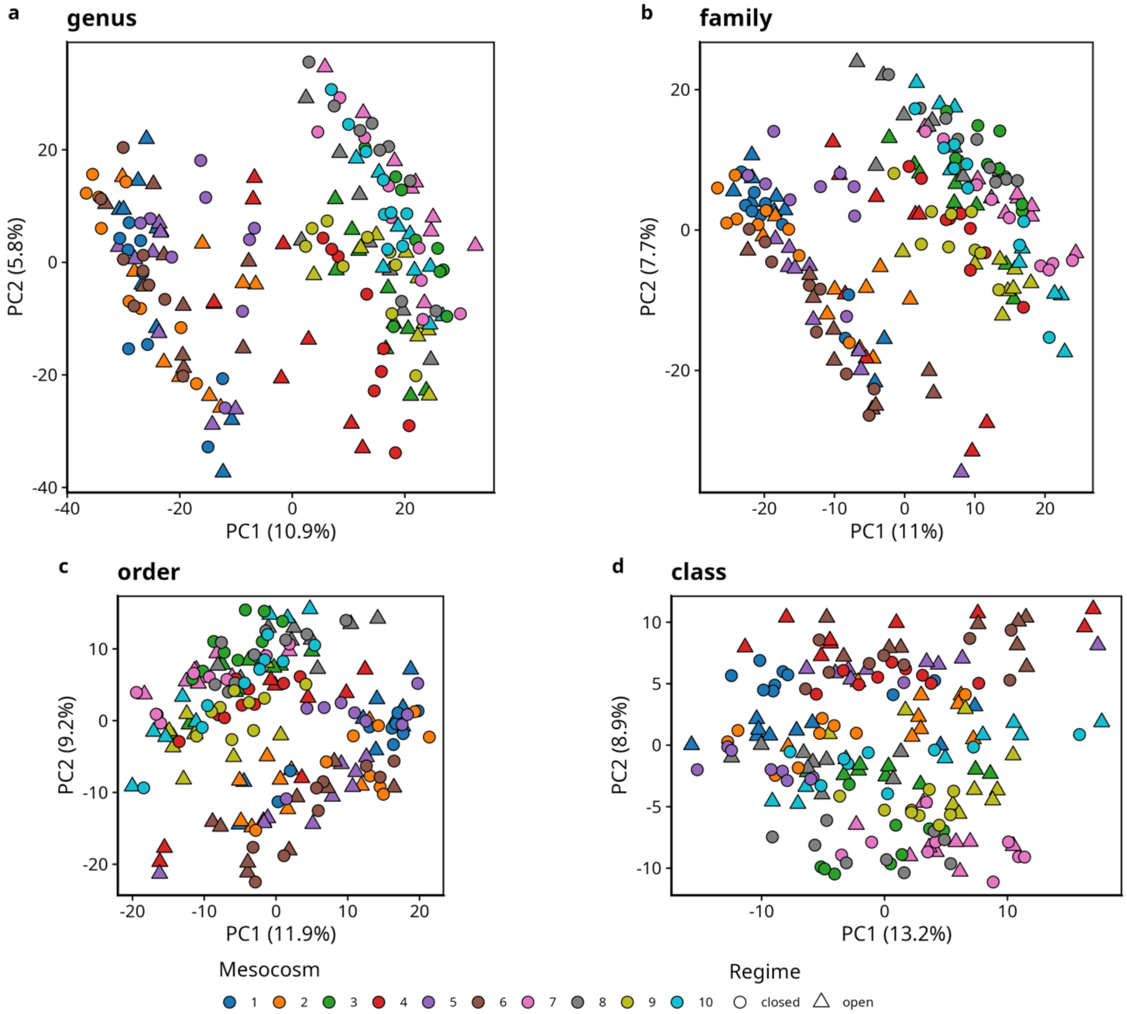


**Figure S7.** Community profiling at higher taxonomic ranks preserves separation into two community types. **a.** Identical to Fig. 1d (main text), included for reference.
**b–d**. PCA based on reads annotated at family (b), order (c), and class (d) ranks, respectively.

**(next page) Figure S8.** Principal Component Analysis (PCA) performed separately for **a.** closed and **b.** open mesocosms, in contrast to the combined PCA in the main text. Community types remain separated along PC1 in both datasets. Ellipses indicate 95% confidence intervals for type A (mesocosms 1, 2, 5, 6) and type B (mesocosms 3, 4, 7–10). **c, d**. Same analyses as (a, b), respectively, but highlighting the top 10 principal components shown in panels (e) and (f). **e,f.** Identities and variance explained by the top 20 components in closed (e) and open (f) mesocosms.

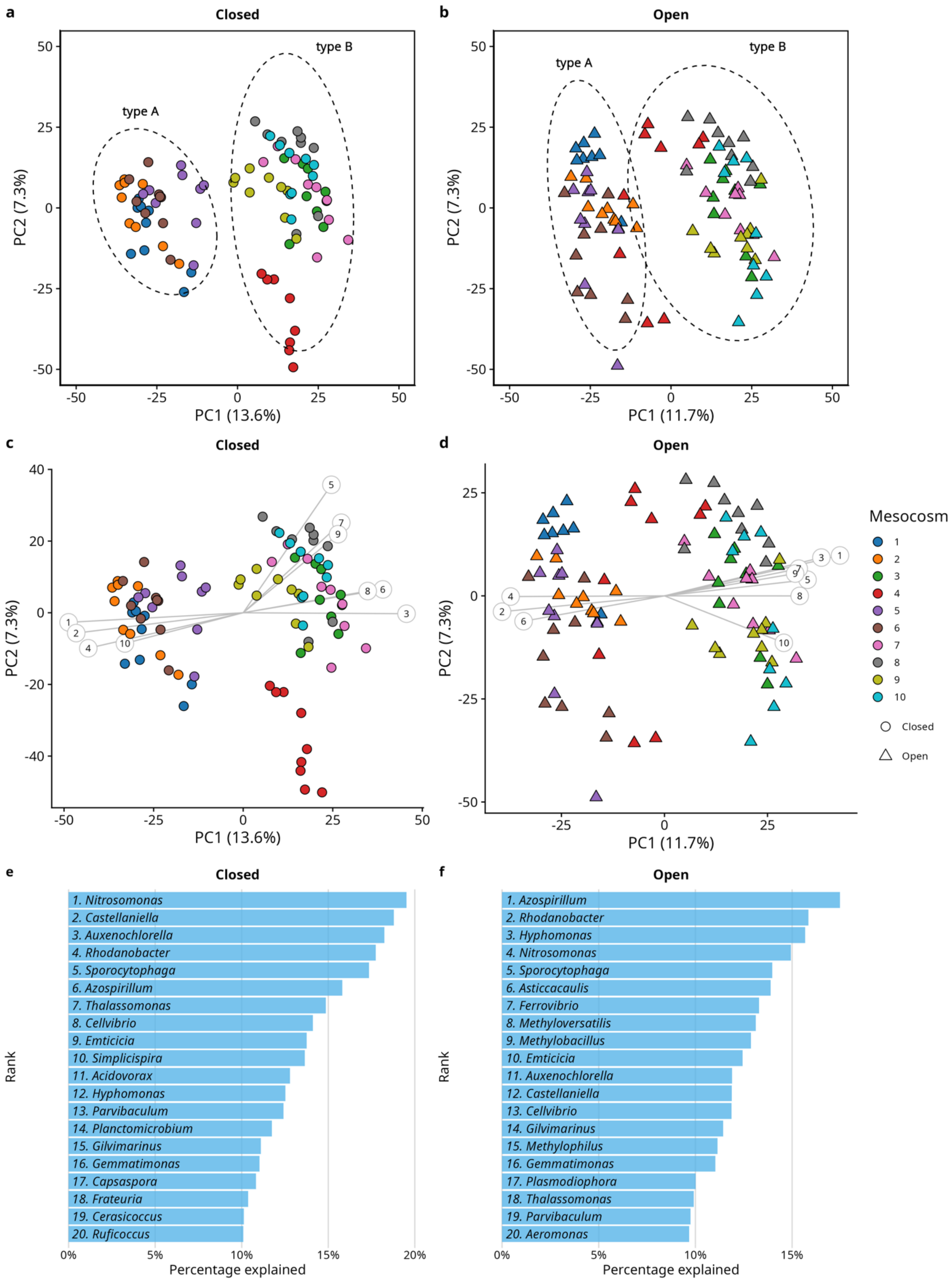

a
Closed
type A
type B
PC1 (13.6%)
PC2 (7.3%)
b
Open
type A
type B
PC1 (11.7%)
PC2 (7.3%)
c
Closed
PC1 (13.6%)
PC2 (7.3%)
d
Open
PC1 (11.7%)
PC2 (7.3%)
Mesocosm
Closed
Open
e
Closed
Rank
1. Nitrosomonas
2. Castellaniella
3. Auxenochlorella
4. Rhodanobacter
5. Sporocytophaga
6. Azospirillum
7. Thalassomonas
8. Cellvibrio
9. Emticicia
10. Simplicispira
11. Acidovorax
12. Hyphomonas
13. Parvibaculum
14. Planctomicrobium
15. Gilvimarinus
16. Gemmatimonas
17. Capsaspora
18. Frateuria
19. Cerasicoccus
20. Ruficoccus
Percentage explained
f
Open
Rank
1. Azospirillum
2. Rhodanobacter
3. Hyphomonas
4. Nitrosomonas
5. Sporocytophaga
6. Asticcacaulis
7. Ferrovibrio
8. Methyloversatilis
9. Methylobacillus
10. Emticicia
11. Auxenochlorella
12. Castellaniella
13. Cellvibrio
14. Gilvimarinus
15. Methylophilus
16. Gemmatimonas
17. Plasmodiophora
18. Thalassomonas
19. Parvibaculum
20. Aeromonas
Percentage explained

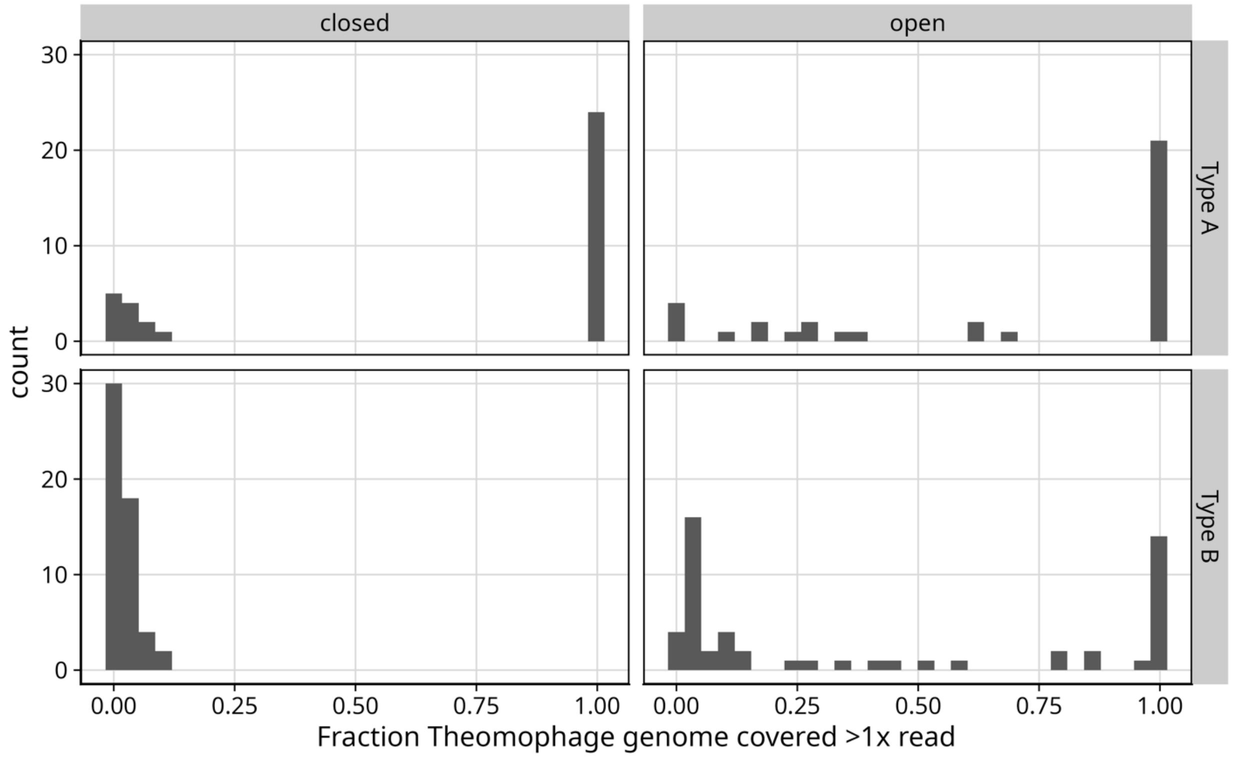


**Figure S9.** Histograms showing the number of samples at a given horizontal coverage (number of nucleotides with ≥1 read mapped) of the Theomophage genome in closed and open samples, split by community type. Low horizontal coverage indicates absence of Theomophage in type-B communities in the closed experimental regime.

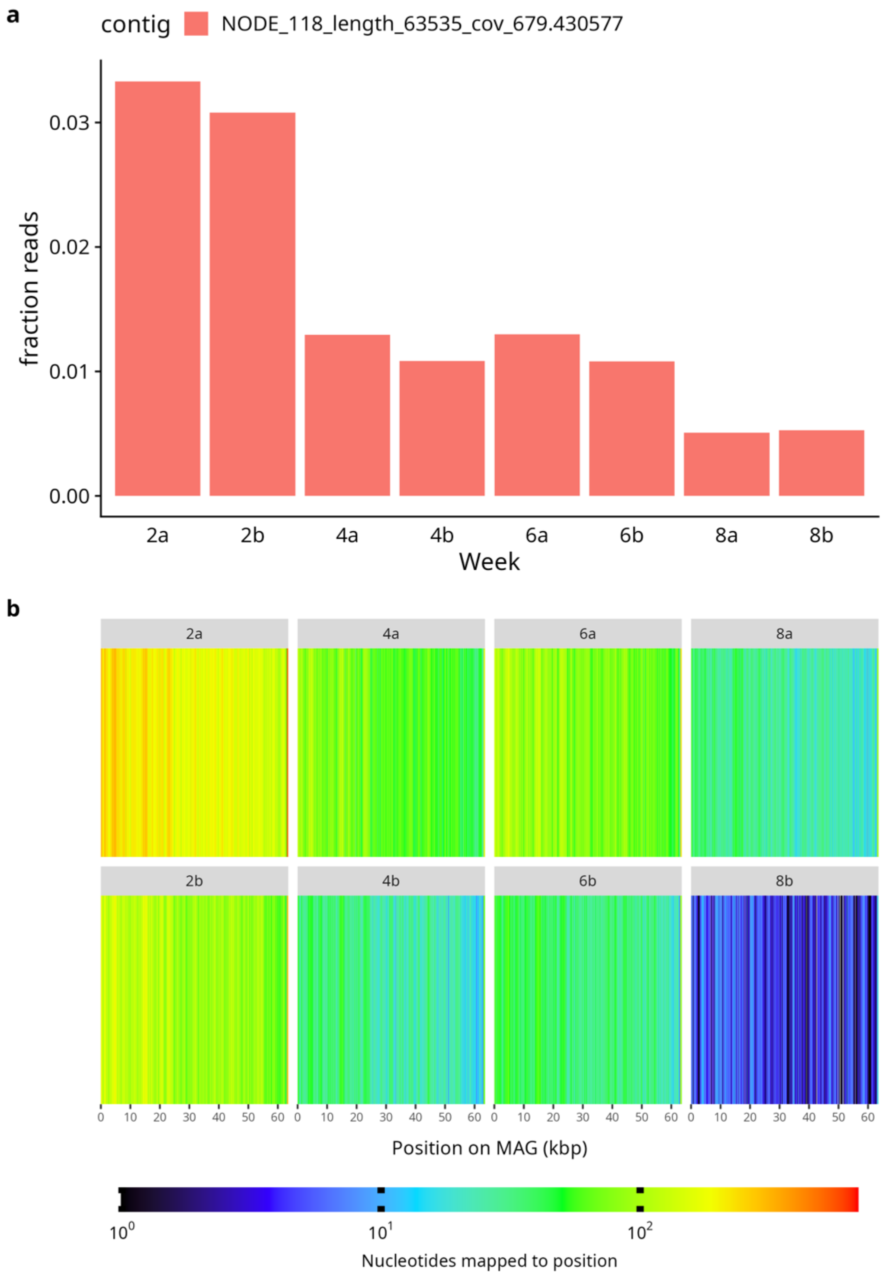


**Figure S10. a.** Relative abundance (fraction of total sample reads) of Theomophage contig NODE_118 in MGE cocktail samples from 0, 2, 4 and 8 weeks. Each sample was sequenced twice, as indicated with letters a,b. **b.** Read coverage profile for the Theomophage MAG confirms detection of the complete genome representation in all samples.

**Figure S11 (next page) Unrooted phylogenetic tree of Terminase large subunit proteins** constructed using five randomly selected representatives per family from the INPHARED dataset, along with the portal protein from Theomophage (see Methods for details). Theomophage clusters with members of the *Schitoviridae* family. Node labels indicate the phage family (by color) and species name.

Terminase large subunit (trimmed alignment)

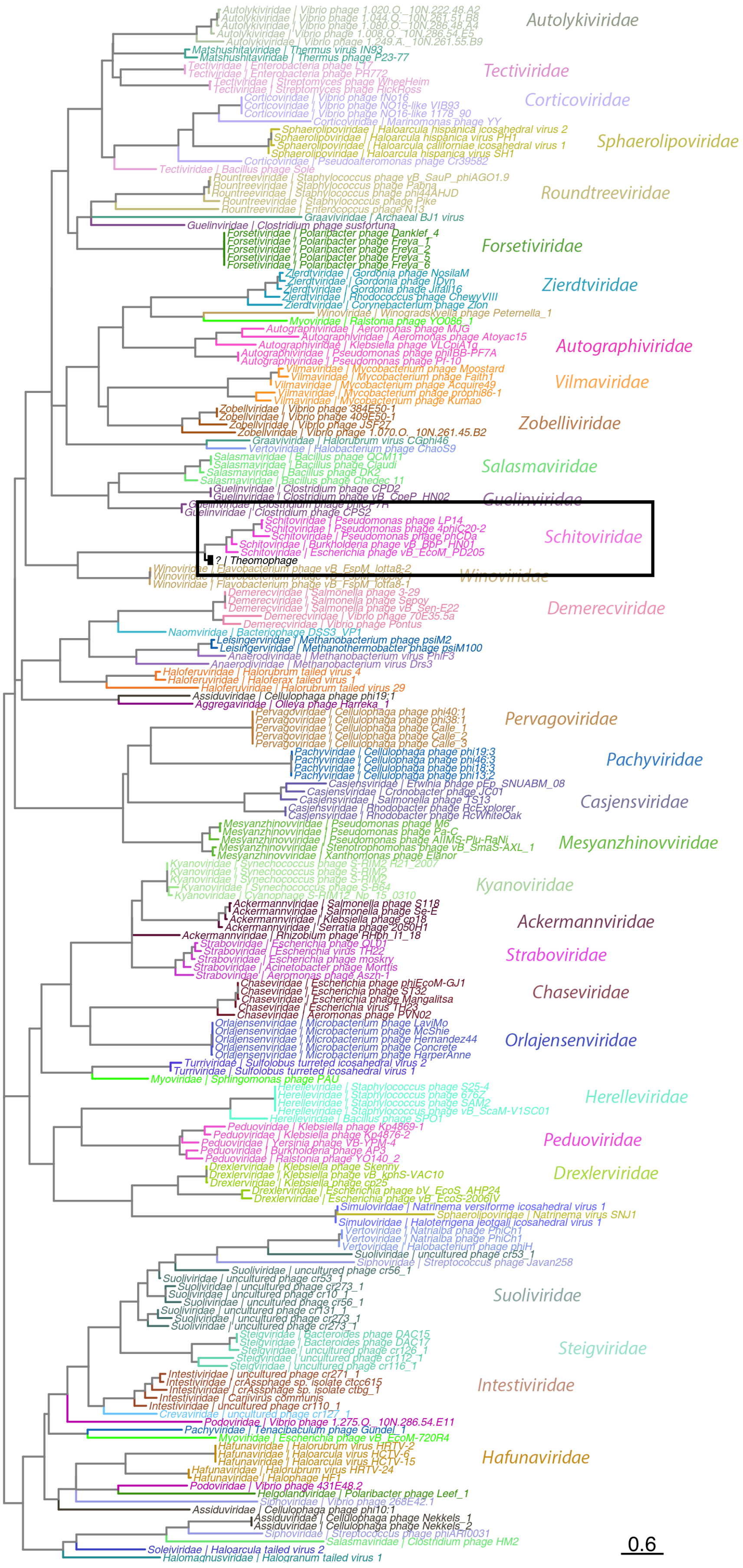

**Figure S12 (next page) Unrooted phylogenetic tree of portal proteins** constructed using five randomly selected representatives per family from the INPHARED dataset, along with the portal protein from Theomophage (see Methods for details). Theomophage clusters with members of the *Schitoviridae* family. Node labels indicate the phage family (by color) and species name.

Portal (trimmed alignment)

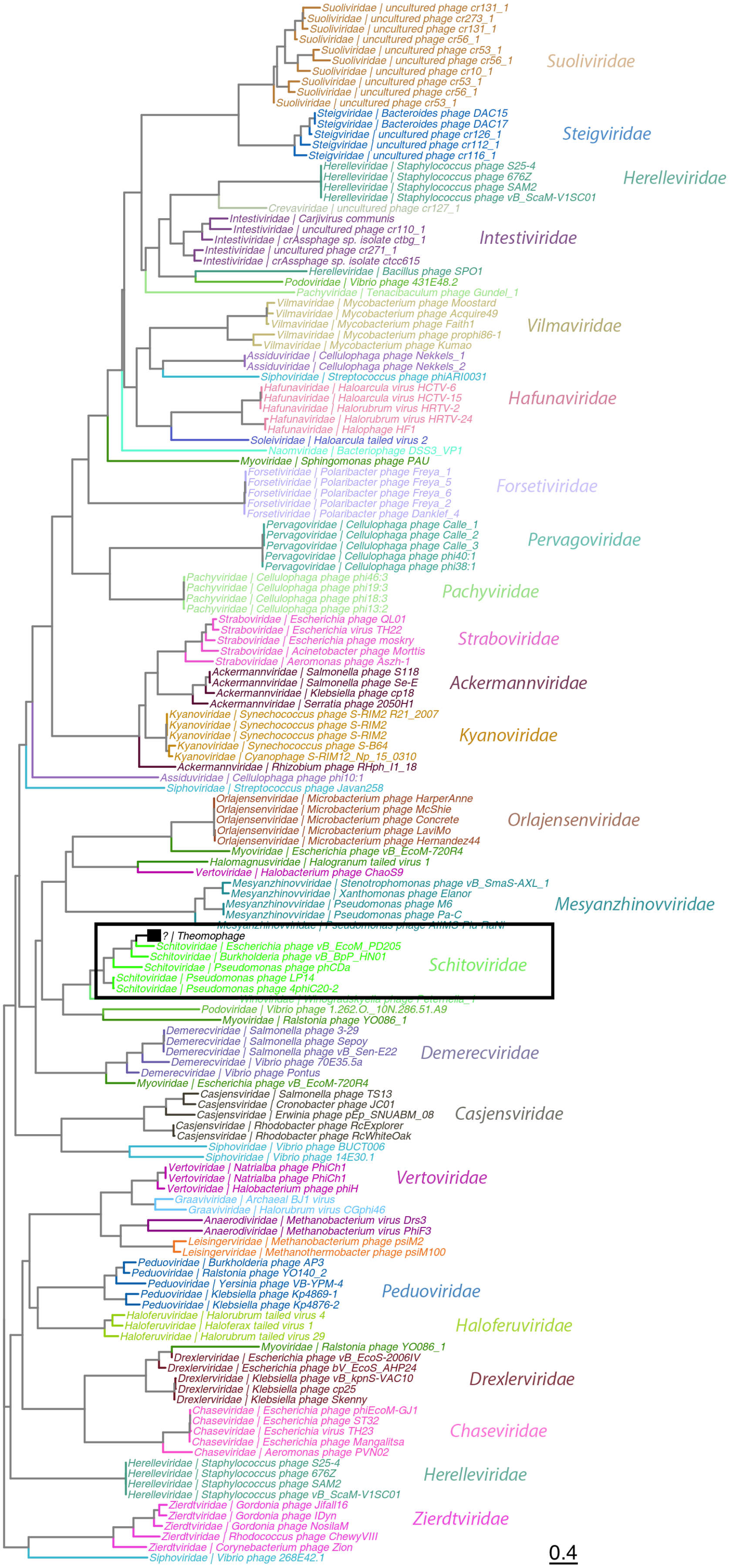

**Figure S13 (next page) Unrooted phylogenetic tree of Major capsid proteins** constructed using five randomly selected representatives per family from the INPHARED dataset, along with the portal protein from Theomophage (see Methods for details). Theomophage clusters with members of the *Schitoviridae* family. Node labels indicate the phage family (by color) and species name.

Major capsid protein (trimmed alignment)

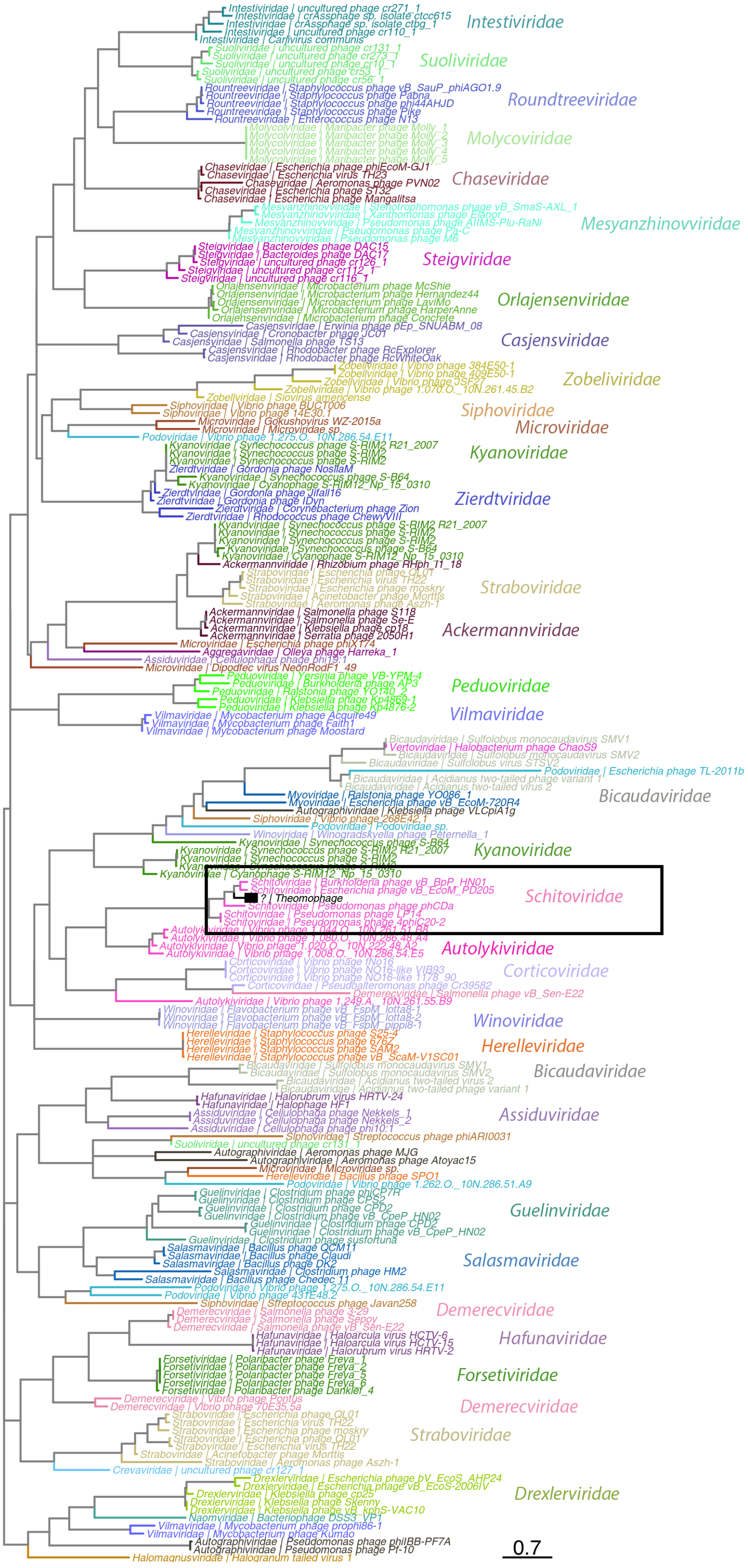

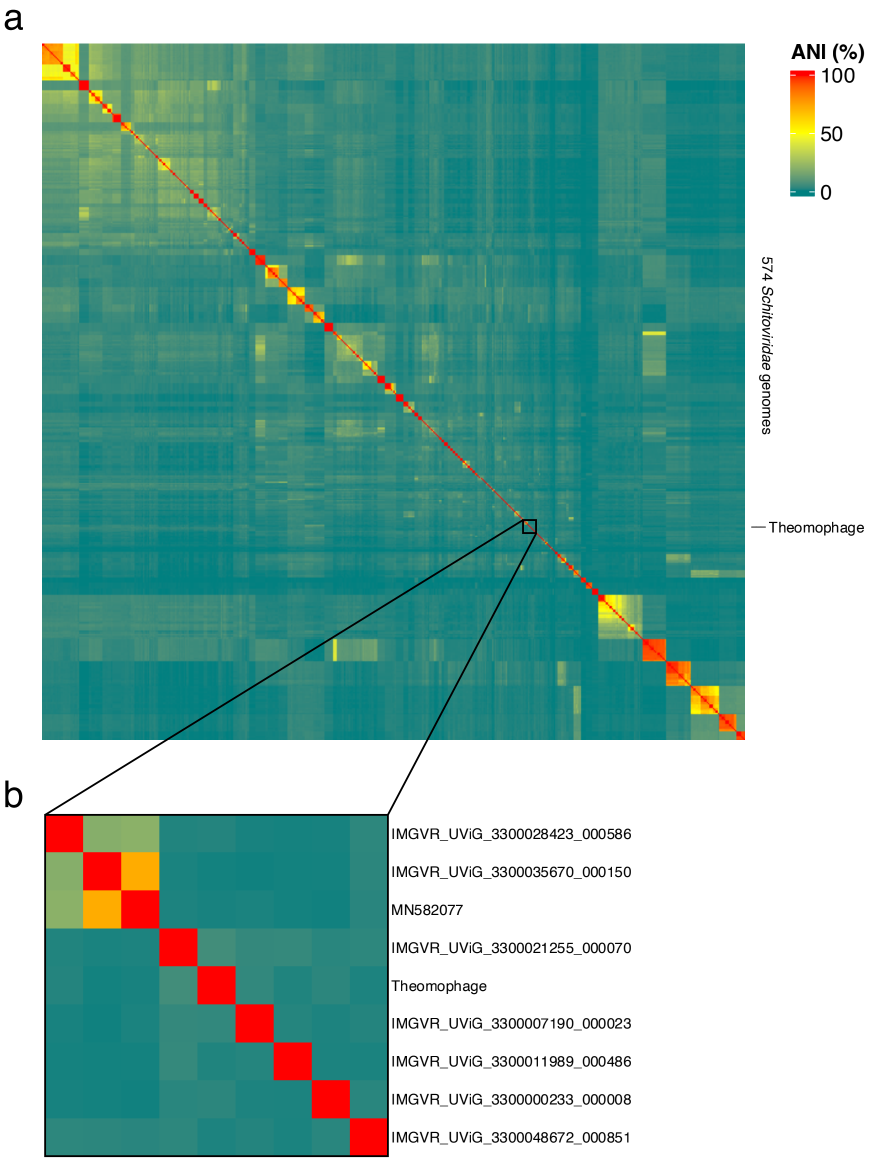


**Figure S14. a.** Heatmap showing VIRIDIC-based pairwise similarity of 573 high-quality (>90% complete) *Schitoviridae* genomes and the Theomophage MAG NODE244. Theomophage shows only 8.98% similarity to known *Schitoviridae*, below the 20% subfamily demarcation threshold, supporting its designation as a novel subfamily. **b.** inset showing location of Theomophage.

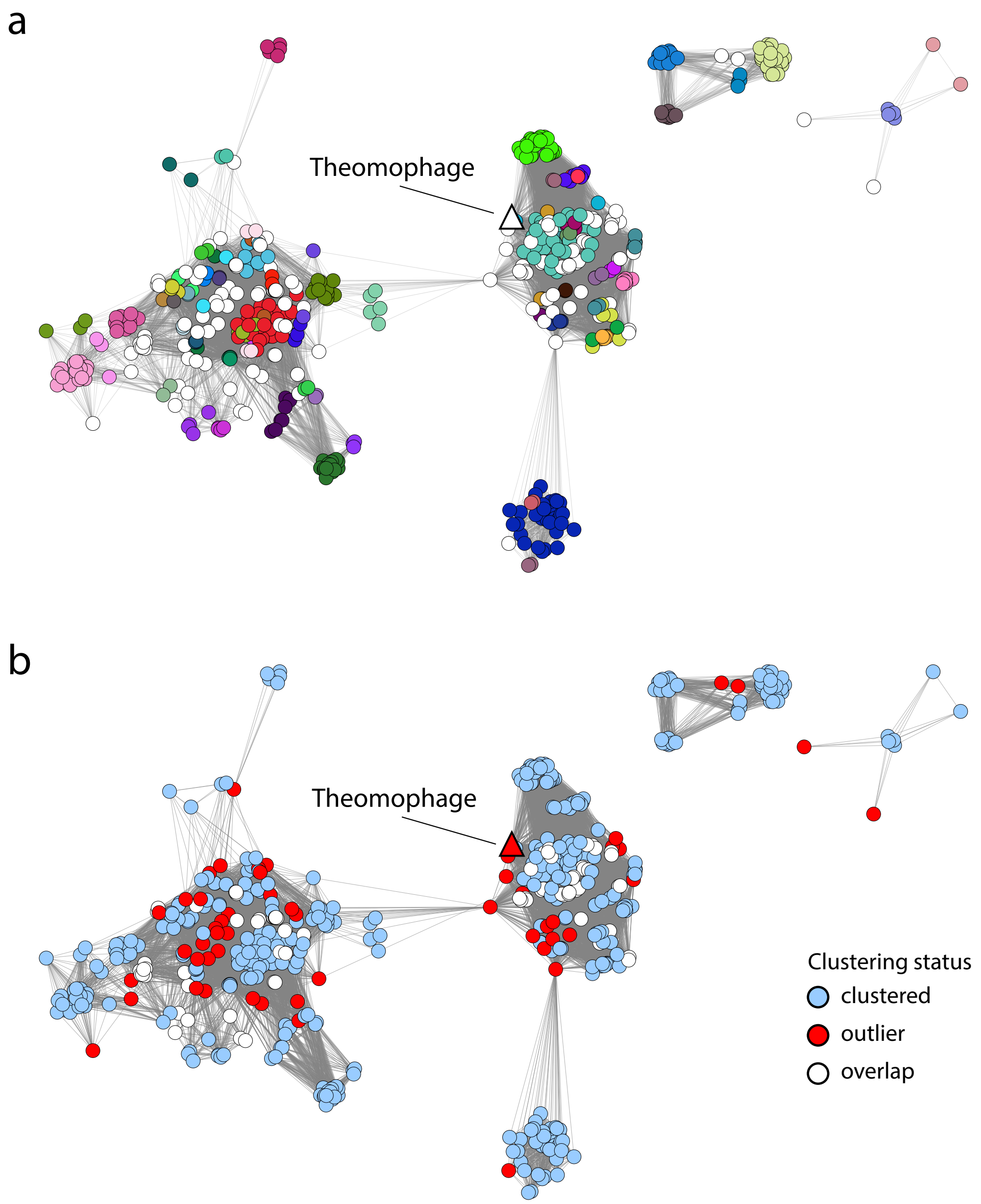


**Figure S15. a.** Gene sharing network of 573 high-quality *Schitoviridae* genomes and Theomophage MAG NODE244, constructed with vConTACT2 using default parameters. Each node represents a viral genome and is colored by the assigned vConTACT2 viral cluster (VC), which reflects ICTV genus level taxonomy. Theomophage (triangle) does not cluster with any known *Schitoviridae* sequences, indicating it is not related at genus level. **b.** Same network as in (a), but nodes colored by clustering status: genomes assigned to a single VC (blue), overlap belonging to multiple VCs (white), and outliers that do not group into any VC (red).

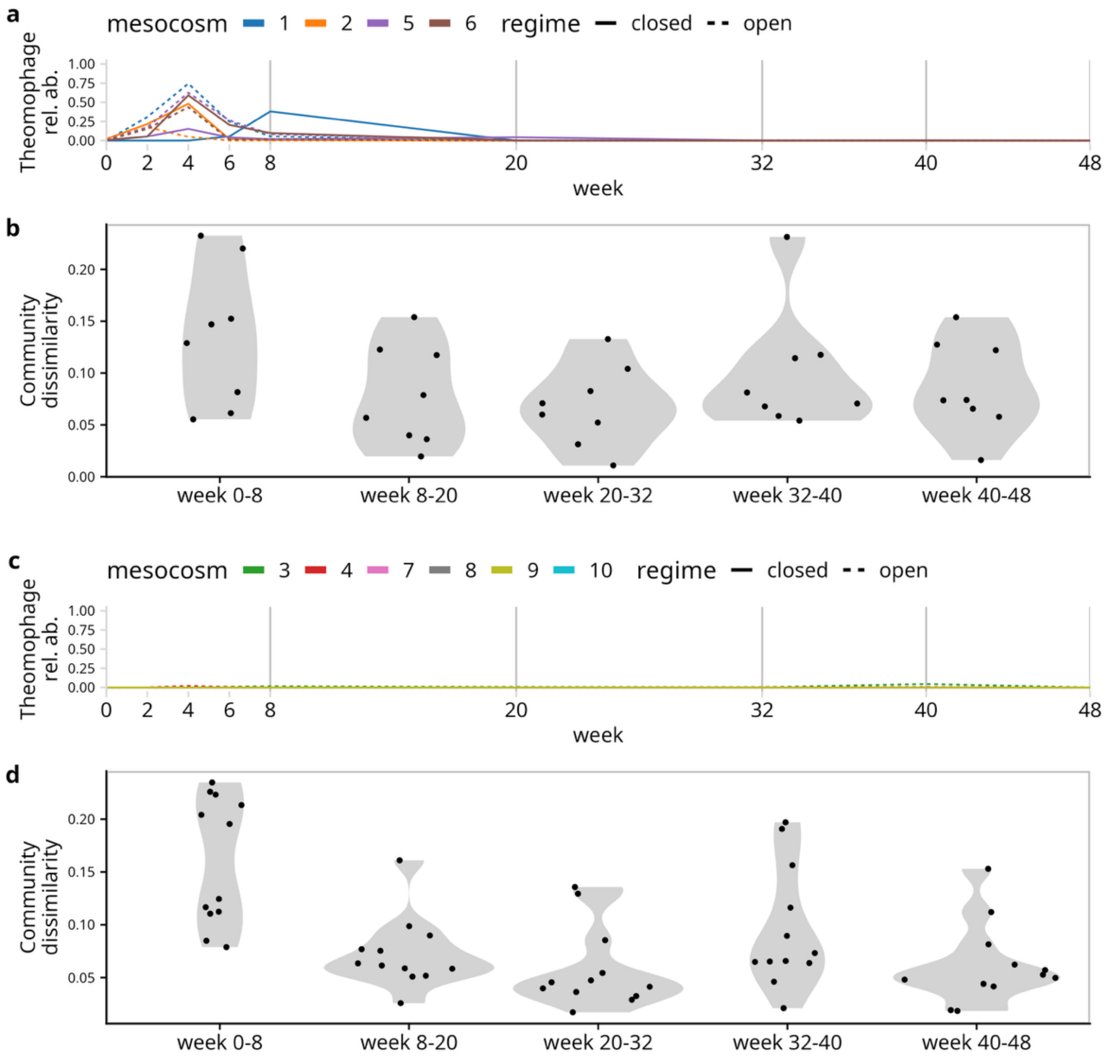


**Figure S16. Theomophage outbreaks do not alter community composition**

**a.** Theomophage abundance in mesocosms with outbreaks (closed and open 1,2,5,6; outbreaks observed between weeks 0-8). **b.** Bray-Curtis dissimilarity of genus abundances for time points spanning Theomophage outbreaks (weeks 0–8) and without outbreaks (all other intervals). While dissimilarities appear higher during outbreaks, complex natural communities typically lose taxa when adapted to lab conditions[1], a trend also observed in mesocosms without outbreaks (d). **c-d** Same as (a-b), but for mesocosms without Theomophage outbreaks. Bray-Curtis dissimilarities were normalized for time interval length.

1. Bittleston, L. S., Gralka, M., Leventhal, G. E., Mizrahi, I. & Cordero, O. X. Context-dependent dynamics lead to the assembly of functionally distinct microbial communities. *Nat. Commun.* **11**, 1440 (2020).

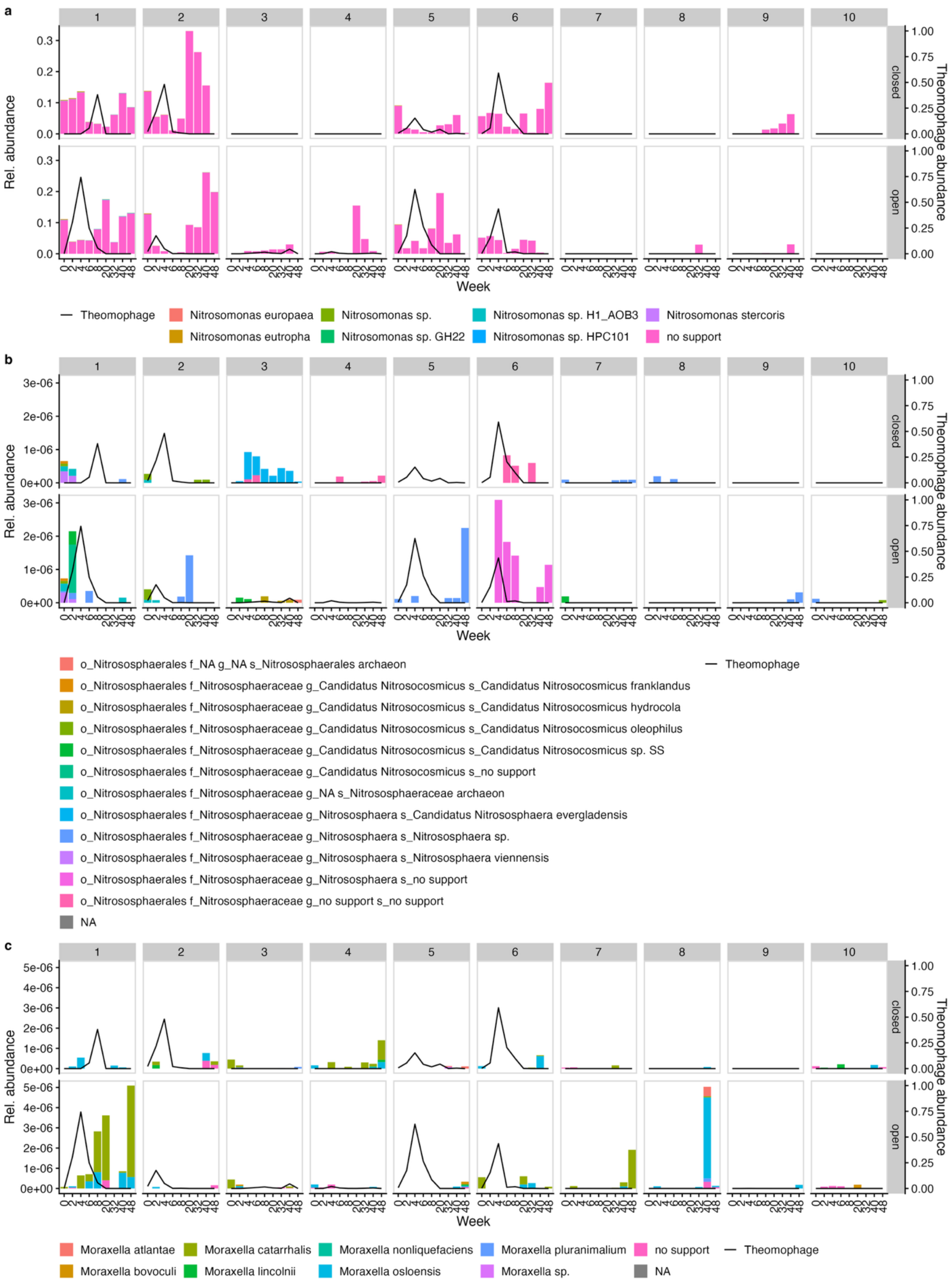
a
1
2
3
4
5
6
7
8
9
10
closed
open
Rel. abundance
Theomophage abundance
Week
Theomophage
Nitrosomonas europaea
Nitrosomonas sp.
Nitrosomonas sp. H1_AOB3
Nitrosomonas stercoris
Nitrosomonas eutropha
Nitrosomonas sp. GH22
Nitrosomonas sp. HPC101
no support
b
Rel. abundance
Theomophage abundance
Week
o_Nitrososphaerales f_NA g_NA s_Nitrososphaerales archaeon
o_Nitrososphaerales f_Nitrososphaeraceae g_Candidatus Nitrosocosmicus s_Candidatus Nitrosocosmicus franklandus
o_Nitrososphaerales f_Nitrososphaeraceae g_Candidatus Nitrosocosmicus s_Candidatus Nitrosocosmicus hydrocola
o_Nitrososphaerales f_Nitrososphaeraceae g_Candidatus Nitrosocosmicus s_Candidatus Nitrosocosmicus oleophilus
o_Nitrososphaerales f_Nitrososphaeraceae g_Candidatus Nitrosocosmicus s_Candidatus Nitrosocosmicus sp. SS
o_Nitrososphaerales f_Nitrososphaeraceae g_Candidatus Nitrosocosmicus s_no support
o_Nitrososphaerales f_Nitrososphaeraceae g_NA s_Nitrososphaeraceae archaeon
o_Nitrososphaerales f_Nitrososphaeraceae g_Nitrososphaera s_Candidatus Nitrososphaera evergladensis
o_Nitrososphaerales f_Nitrososphaeraceae g_Nitrososphaera s_Nitrososphaera sp.
o_Nitrososphaerales f_Nitrososphaeraceae g_Nitrososphaera s_Nitrososphaera viennensis
o_Nitrososphaerales f_Nitrososphaeraceae g_Nitrososphaera s_no support
o_Nitrososphaerales f_Nitrososphaeraceae g_no support s_no support
NA
Theomophage
c
Rel. abundance
Theomophage abundance
Week
Moraxella atlantae
Moraxella catarrhalis
Moraxella nonliquefaciens
Moraxella pluranimalium
no support
Theomophage
Moraxella bovoculi
Moraxella lincolnii
Moraxella osloensis
Moraxella sp.
NA

**Figure S17 (previous page).** Abundance of putative Theomophage hosts across 10 closed and open compost mesocosms. Host predictions were made using CRISPR spacer matching against the SpacerDB global spacer database and genome-based host prediction via iPHoP (see Methods and **Table S8-S10**). Theomophage abundance is shown as a black line (secondary y-axis).

**a.** *Nitrosomonas* species, identified as a candidate host based on six spacer matches with 2-3 mismatches in SpacerDB, providing weak evidence of a host relationship. However, *Nitrosomonas* was consistently associated with type-A communities where Theomophage outbreaks occurred (see Fig.1b-e) and was detected in 2 of the 5 mesocosms where Theomophage invaded (open_3 and open_4). Note that no CRISPR spacers targeting Theomophage were found in *Nitrosomonas* sequences from the compost metagenomes.

**b.** *Moraxella* species. A single spacer with one mismatch was found in SpacerDB, originating from a wastewater-associated *Moraxella* sequence.

**c.** Taxa within the *Nitrososphaerales* order, predicted as a candidate host by iPHoP.

Both *Moraxella* and *Nitrososphaerales* were detected at very low abundance (0.000509% and 0.000309% of sample reads, respectively), making them unlikely hosts for the abundant Theomophage. However, since metagenomes were collected at the end of a two-week growth cycle, it remains possible that active phage predation led to host population collapse, reducing their population sizes to near the detection threshold at the time of sampling.

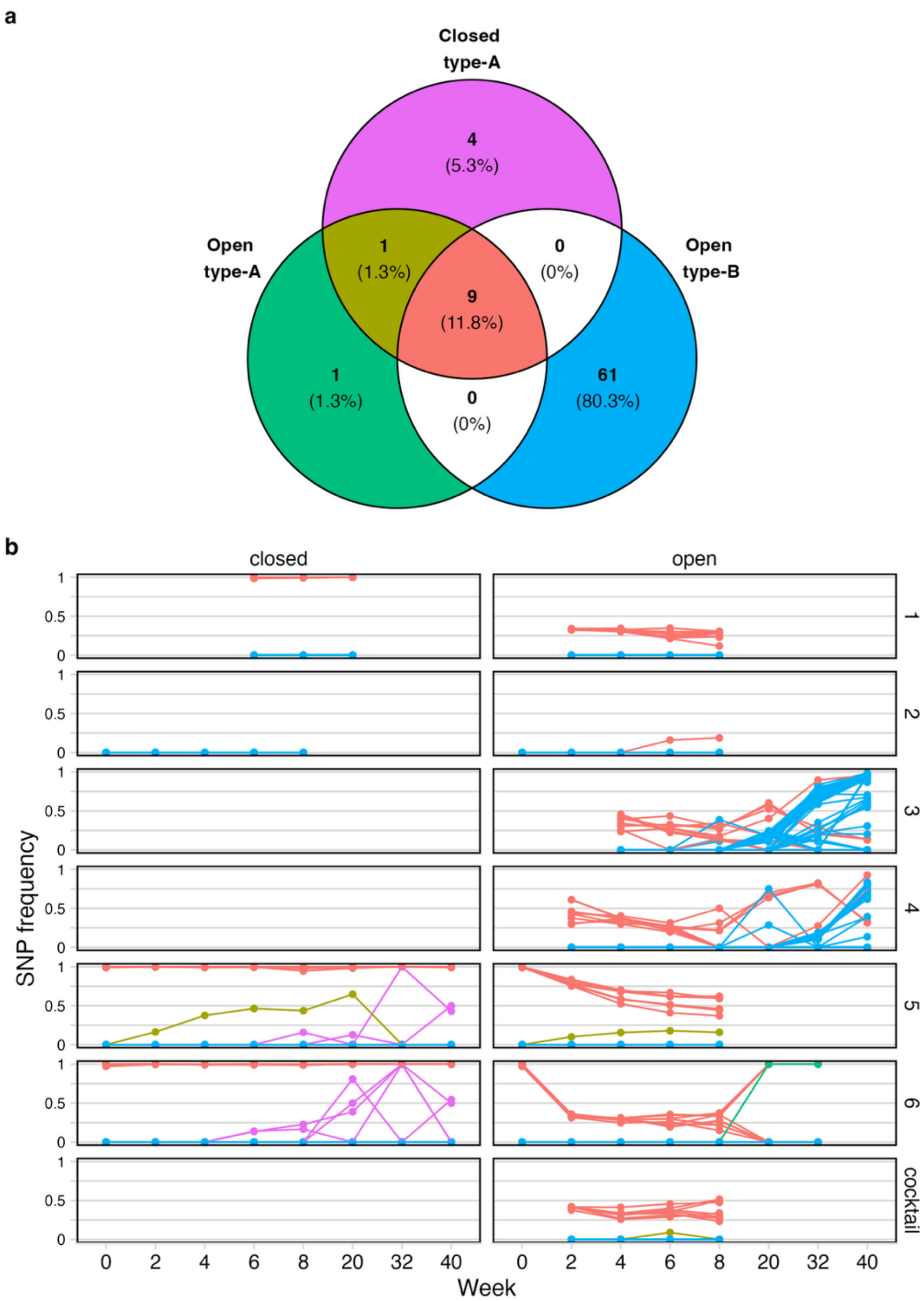


**Figure S18. a.** Venn diagram of 76 Theomophage SNPs detected across native (type-A) and novel (type-B) communities in closed and open regimes. Theomophage was not detected in type-B communities in closed mesocosms. **b.** Allele frequency trajectories, with colors corresponding to (a). Olive and pink SNPs represent microdiversity within genotype G2 in mesocosms closed_5, closed_6 and open_5. The olive SNP is unique to community 5 and observed in both regimes. The green SNP is unique to open_6.

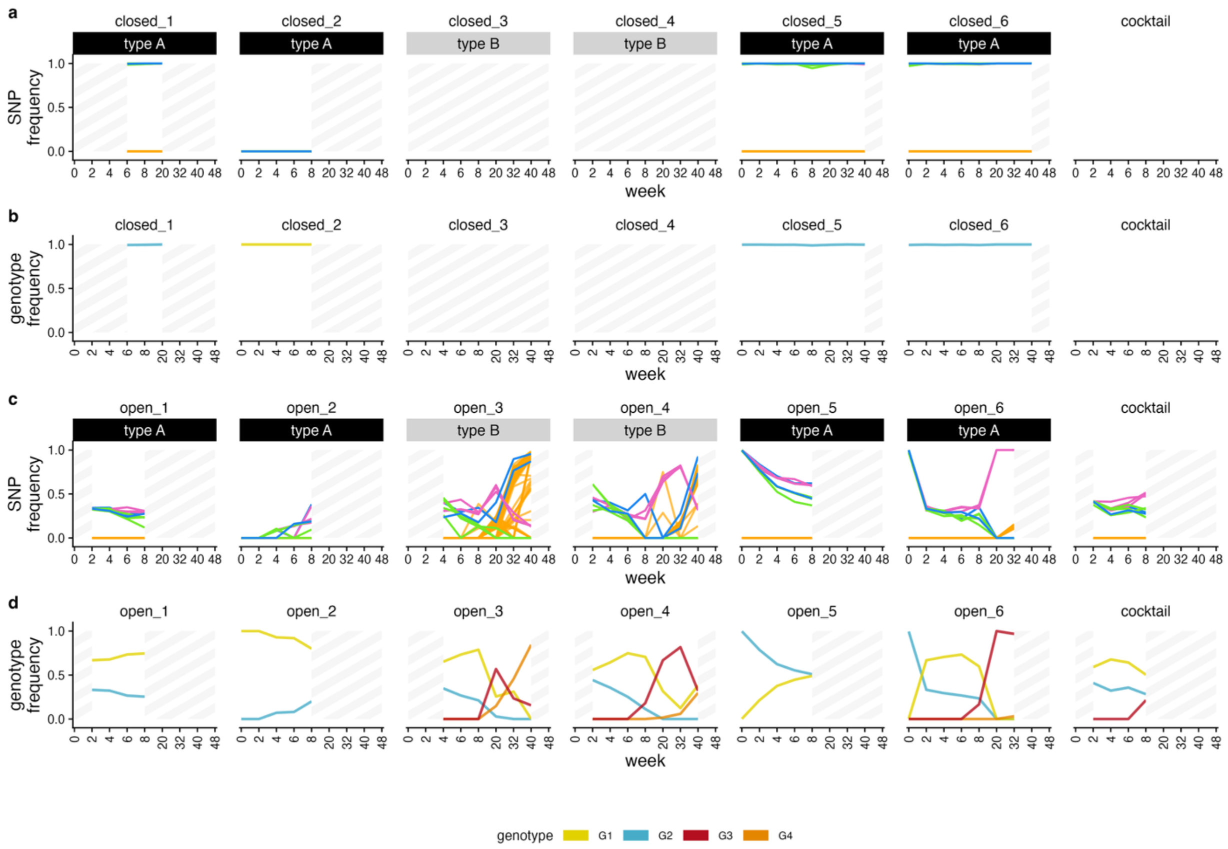


**Figure S19. More sensitive SNP analysis reveals additional migration events between open mesocosms.** Using the same approach as in Figure 4 of the main text, but with less strict variant filtering reveals that all nine SNPs defining genotype G2 were present at week 8 in mesocosm open_2. Genotype G4 was also detected in open_6 at week 32, indicating migration to this mesocosm. SNP and genotype colors match Fig.4 and Table 1 of the main text.

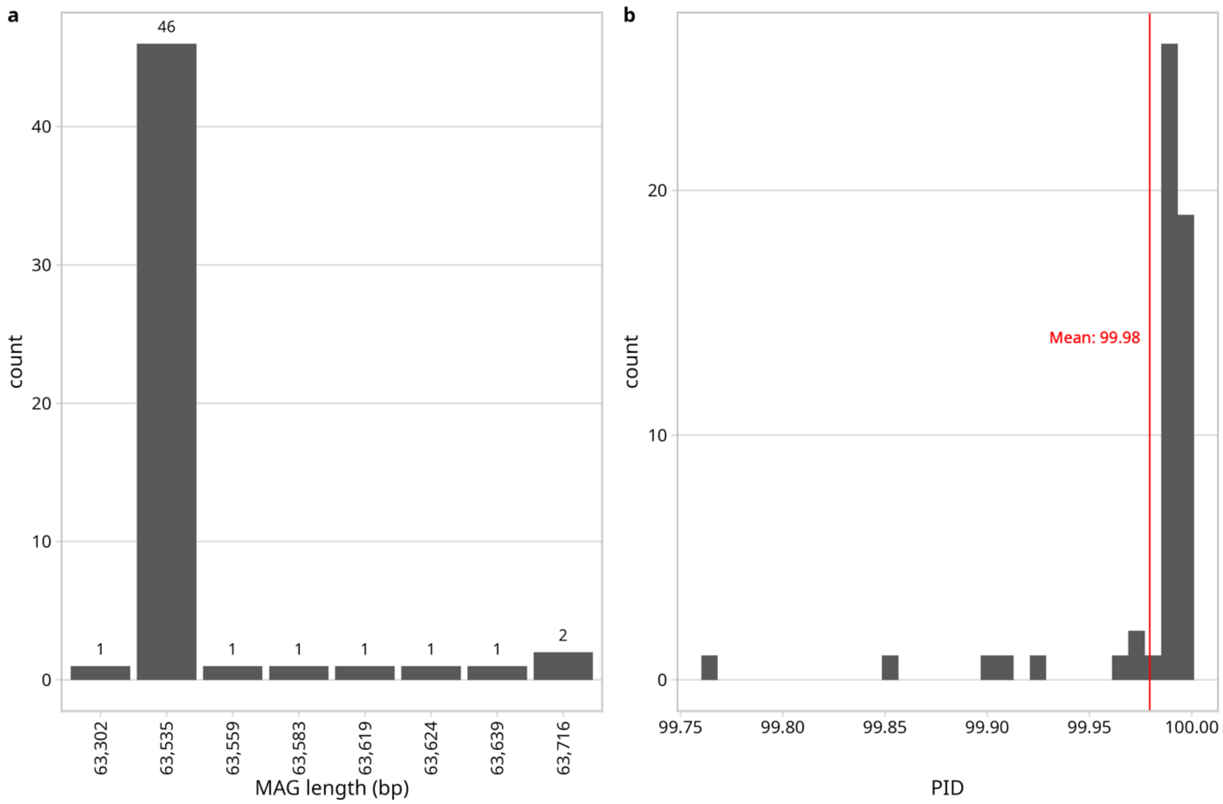


**Figure S20. a.** Length distribution of Theomophage contigs >60kb from 170 separately assembled metagenomes. Length of assembled contigs was highly consistent. Alignment and read pileup inspection (not shown) indicated contigs >63,535 bp are chimeric assemblies of multiple Theomophage genotypes. **b.** Sequence similarity of contigs from (a), calculated with VIRIDIC (see Methods).

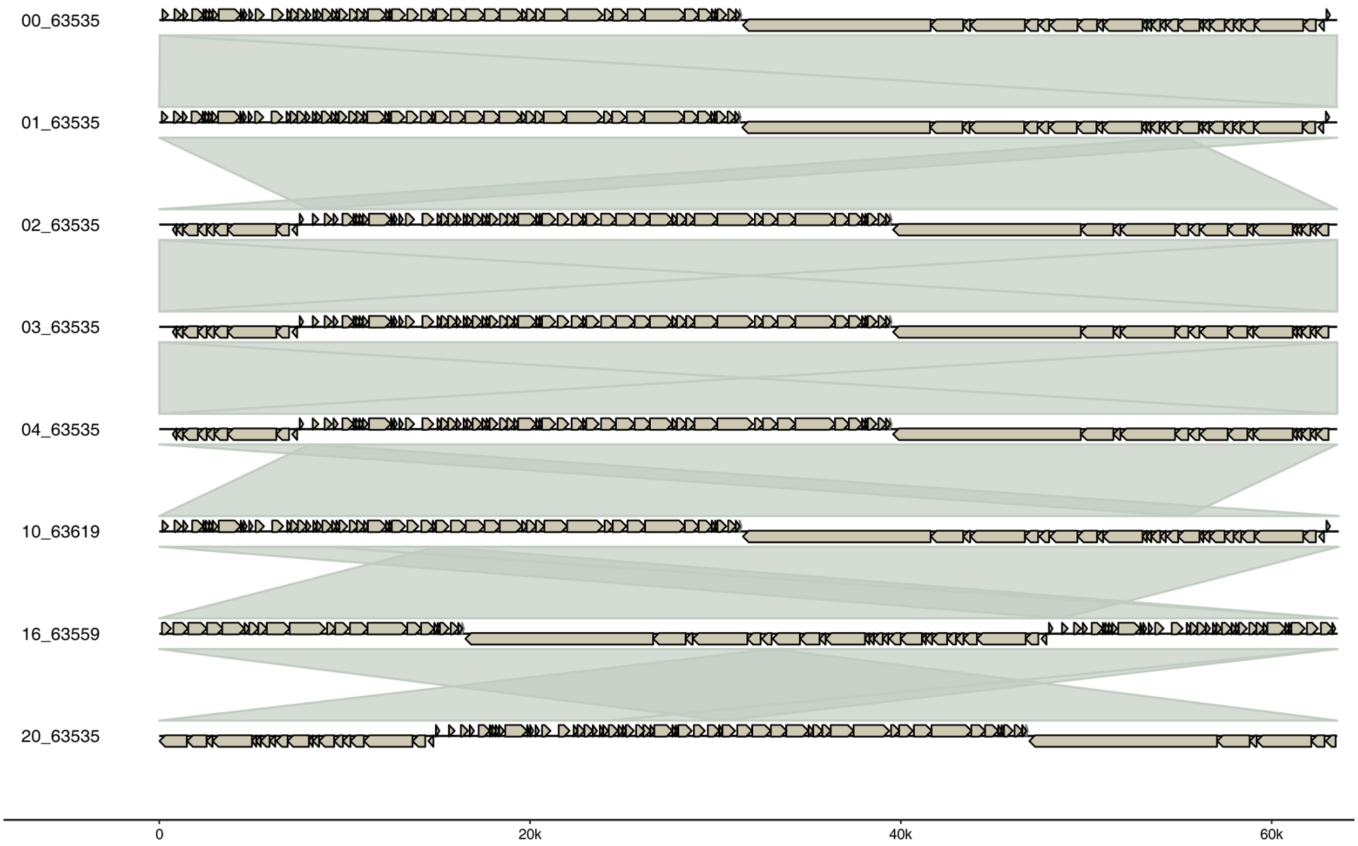


**Figure S21.** Theomophage contigs independently assembled from closed_5 samples from transfer 0, 1, 2, 3, 4, 10, 16, and 20. Contigs are circularly permuted relative to each other and have highly consistent length, indicating complete assembly of a bacteriophage genome. Labels indicate transfer number and contig length.
